# A pair of effectors encoded on a conditionally dispensable chromosome of *Fusarium oxysporum* suppress host-specific immunity

**DOI:** 10.1101/2020.10.06.329052

**Authors:** Yu Ayukawa, Shuta Asai, Pamela Gan, Ayako Tsushima, Yasunori Ichihashi, Arisa Shibata, Ken Komatsu, Petra M. Houterman, Martijn Rep, Ken Shirasu, Tsutomu Arie

## Abstract

Many plant pathogenic fungi contain conditionally dispensable (CD) chromosomes that are associated with virulence, but not growth *in vitro*. Virulence-associated CD chromosomes carry genes encoding effectors and/or host-specific toxin biosynthesis enzymes that may contribute significantly to determining host specificity. *Fusarium oxysporum* causes devastating diseases of more than 100 plant species. Among a large number of host-specific forms, *F. oxysporum* f. sp. *conglutinans* (*Focn*) can infect Brassicaceae plants including Arabidopsis and cabbage. Here we show that *Focn* has multiple CD chromosomes involved in virulence but also in vegetative growth, which is an atypical feature of CD chromosomes. We identified specific CD chromosomes that are required for virulence on Arabidopsis, cabbage, or both, and describe a pair of effectors encoded on one of the CD chromosomes that is required for suppression of Arabidopsis-specific phytoalexin-based immunity. The effector pair is highly conserved in *F. oxysporum* isolates capable of infecting Arabidopsis, but not of other plants. This study provides insight into how host specificity of *F. oxysporum* may be determined by a pair of effector genes on a transmissible CD chromosome.

## Introduction

Pathogenic fungi often carry chromosomes that are not necessary for growth in the non-pathogenic state^1, 2^. Analogous to the well-characterized virulence plasmids in bacteria, the number of these dispensable chromosomes in individual isolates can vary. In plant pathogenic fungi, dispensable chromosomes that are associated with virulence are generally referred to as supernumerary, ‘B’, or conditionally dispensable (CD) chromosomes^1^. When pathogenic fungi lack CD chromosomes, they can grow *in vitro*, but often exhibit attenuated or no virulence^1, 2^. The functions of CD chromosomes in some plant pathogenic fungi are associated with suppression or deactivation of host-specific factors. In *Fusarium solani*, for example, a CD chromosome carries phytoalexin detoxifying genes^3^. In contrast, the CD chromosomes of *Alternaria alternata* and *Cochliobolus carbonum* harbor host-specific toxin genes^4^. Therefore, CD chromosomes can be crucial determinants of host specificity that are defined by phytotoxin activity or by defense against chemicals such as phytoalexins.

*Fusarium oxysporum* causes devastating diseases of more than 100 plant species, including economically important crops such as tomato, banana and melon^5^. Individual isolates of *F. oxysporum* have different host ranges and are classified into *formae speciales* (ff. spp.) based on the susceptibility of plant species to infection. Although much is known about the genetics and pathology of *F. oxysporum*, the precise molecular mechanisms of host specificity remain unclear. So far, CD chromosomes have been identified in the tomato-infecting pathogen *F. oxysporum* f. sp. *lycopersici* (*Fol*) and in *F. oxysporum* f. sp. *radicis-cucumerinum* (*Forc*), a cucurbit-infecting pathogen^6–8^. The *Fol* isolate 4287 and the *Forc* isolate 016 each contain a single virulence-associated CD chromosome that is transferable to other isolates^7–9^. Horizontal transfer of the CD chromosomes from *Fol*4287 or *Forc*016 converts non-pathogenic *F. oxysporum* isolates into pathogens of their respective hosts^6, 7, 9^. Part of this phytopathogenic conversion is often due to the expression of CD-encoded effectors that modulate host immunity against infection, such as Secreted In Xylem (SIX) effectors that are, as their name indicates, secreted into xylem elements during infection^10, 11^. A total of fourteen *SIX* genes (*SIX1* to *14*) have been identified from *Fol*^10^. The CD chromosome of *Fol*4287 contains all of the *SIX* genes except *SIX4*, which is not present in *Fol*4287^6, 12^, but is present in certain other *Fol* isolates. The CD chromosome of *Forc*016 contains *SIX6*, *SIX9*, *SIX11* and *SIX13* homologs^7^. *SIX1*, *SIX3*, *SIX5* and *SIX6* from *Fol*, and *Forc*016 *SIX6* are involved in overcoming resistance in tomato and cucumber, respectively^7, 10^, but their molecular mechanisms as virulence factors are as yet unknown.

Arabidopsis-infecting isolates of *F. oxysporum* are useful as a model pathosystem. There are at least three ff. spp. that cause disease on Arabidopsis: f. sp. *conglutinans*, f. sp. *matthiolae* and f. sp. *raphani*^13^. *F. oxysporum* f. sp. *conglutinans* (*Focn*) can also infect other Brassicaceae plants such as cabbage (*Brassica oleracea*). The *SIX1* gene is required for full virulence on cabbage in *Focn*^14^, but the *Focn* factor(s) that are required for virulence on Arabidopsis have not been identified. We have previously shown that the *Focn* isolate Cong:1-1 (*Focn*Cong:1-1) harbors *SIX1*, *SIX4*, *SIX8* and *SIX9* homologs on multiple chromosomes of different sizes^15^. Although these chromosomes are presumed to be conditionally dispensable in *Focn*, their status as CD chromosomes has not been established.

Here we report, through analyses of chromosome-deficient *Focn*Cong:1-1 strains and through horizontal chromosome transfer, that *Focn*Cong:1-1 has multiple atypical CD chromosomes involved in both vegetative growth and virulence. Importantly, we identified individual CD chromosomes that are required for virulence on Arabidopsis, cabbage, or both. Furthermore, we identified a pair of effector genes on a CD chromosome that are required for suppression of Arabidopsis-specific phytoalexin-based immunity.

## Results

### Chromosome-level genome assembly of *Focn*Cong:1-1

We assembled the *Focn*Cong:1-1 genome sequence into 198 contigs with an N50 of 1.271 Mb. To improve contiguity, we further performed optical mapping using two restriction enzymes. The final assembly consisted of 22 scaffolds (SCs) with an N50 SC length of 4,865 kb and a 99.1% complete BUSCO score (Table 1). For gene prediction, we generated transcriptome data from axenic culture and plant infections, resulting in a total of 21,781 genes, among which are eight presumptive effector genes (*SIX1*, *SIX4*, *SIX8*, *SIX9* and *FOA1*-*FOA4*) that were previously known from Arabidopsis-infecting *F. oxysporum*^16^, as well as the homologous genes of *FOA1* and *FOA4,* which were named *FOA1b* and *FOA4b*, respectively. To find unknown effectors, 1,467 putative secreted proteins were screened for proteins with an effector-like structure using the EffectorP v1 and/or v2 algorithm^17, 18^. A total of 263 secreted proteins were predicted as effectors by both EffectorP v1 and v2. This prediction did not include FOA1, which is involved in the suppression of pattern-triggered immunity^16^, nor its homolog FOA1b. Therefore, a total of 265 proteins, including FOA1 and FOA1b, were defined as high-confidence effector candidates (Table 1 and Supplementary Data 1).

**Table 1.** *Focn* Cong:1-1 genome statistics.

| Statistics | Value |
| --- | --- |
| Assembly stats |  |
| Assembly size (Mb) | 68.8 (72.2)<sup>a</sup> |
| No. of scaffolds | 22 |
| No. of non-scaffolded contigs | 125 |
| Max scaffold length (kb) | 7,006 |
| N50 scaffold length (kb) | 4,865 |
| GC content (%) | 48.40 |
| BUSCO coverage (%) | 99.1 |
| Gene models |  |
| Total no. of genes | 21,781 |
| Total no. of proteins | 22,094 |
| No. of genes encoding secreted proteins | 1,467 |
| No. of high-confidence effector candidates: | 265<sup>b</sup> |
| Predicted by both EffectorP v1 and v2 | 263 |
| Predicted by either EffectorP v1 or v2 | 393 |

<sup>a</sup>The number in parentheses is total assembly size including non-scaffolded contigs.
<sup>b</sup>FOA1 and FOA1b are added to the list of effector candidates predicted by both EffectorP v1 and v2.

The *F. oxysporum* genome is composed of core genomic regions that are conserved among *Fusarium* species, and additional accessory genomic regions that are conserved in certain isolates^19^. Comparative analysis with the *Fol*4287 genome as a reference indicated that (i) the *Focn*Cong:1-1 SCs have no homology with known accessory genomic regions in *Fol*4287 (chr01B; chr02B; chr03; chr06; chr14; chr15)^6^, (ii) similarly, there are genomic regions of *Focn*Cong:1-1 that have no homology with *Fol*4287, and (iii) the non-homologous genomic regions are enriched in transposable elements (TEs) (Fig. 1a). All known effector genes except *FOA4* are located in the TE-rich genomic region in *Focn*Cong:1-1 as follows: *SIX1* (in SC8), *SIX4* (SC9), *SIX8* (SC10), *SIX9* (SC3), *FOA1* (SC5), *FOA1b* (SC10), *FOA2* (SC9), *FOA3* (SC3), and *FOA4b* (SC10). *FOA4* (SC12) may be a pseudogene since it is not expressed either *in vitro* or *in planta* (Supplementary Data 2). TEs are suspected to be involved in the generation of genomic variations leading to environmental adaptation and, in the case of pathogens, they may have been involved in the acquisition of the ability to infect particular hosts^20^. Therefore, chromosomes containing TE-enriched genomic regions have a high potential to be CD chromosomes.

**Figure 1.**
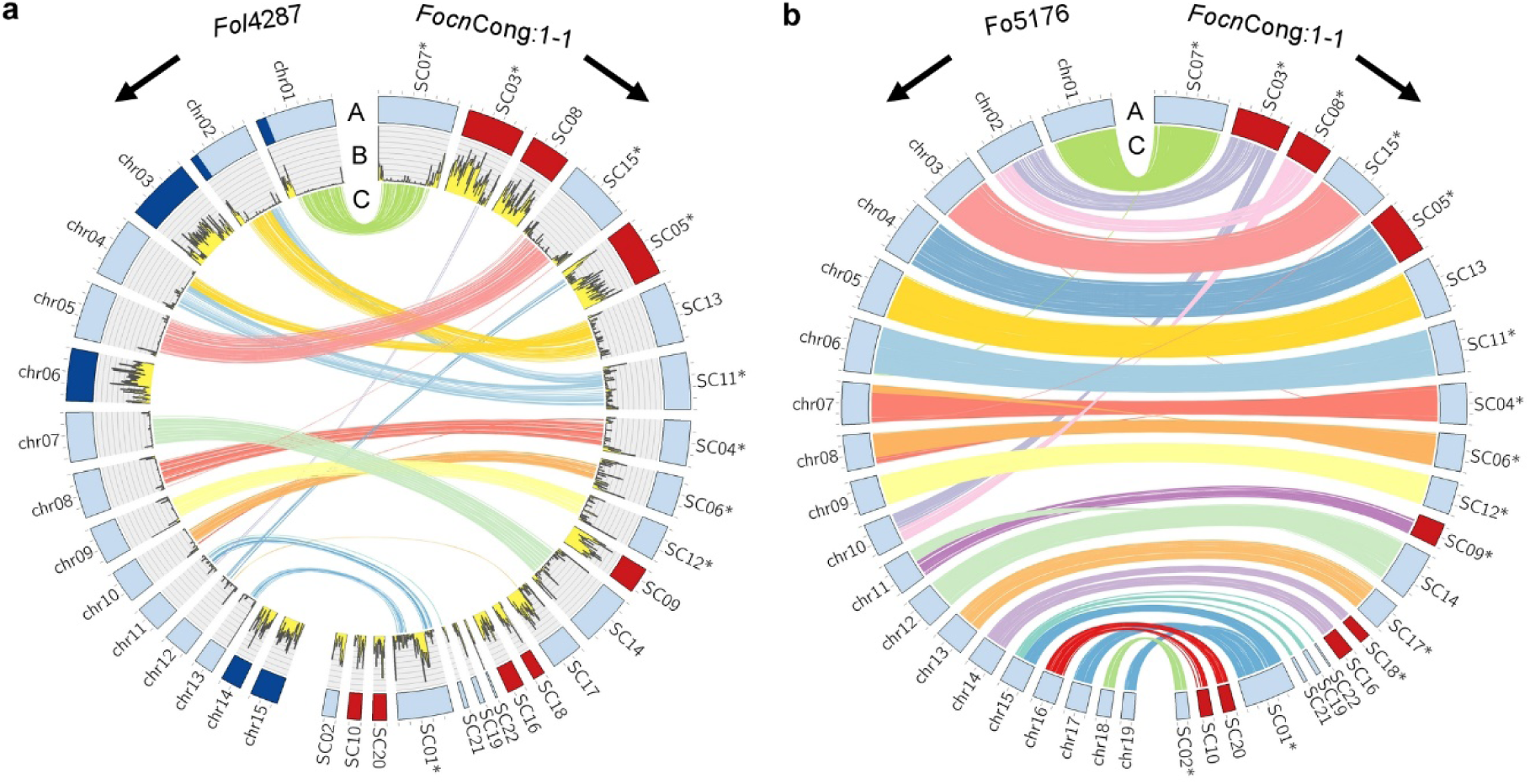
Comparison of whole genome assemblies among *Focn*Cong:1-1, *Fol*4287 and Fo5176. Whole genome assemblies were compared between *Fol*4287 and *Focn*Cong:1-1 (**a**) and between Fo5176 and *Focn*Cong:1-1 (**b**). Ring A: Circular representation of the pseudomolecules. Red and dark blue indicate dispensable genomic regions in *Focn*Cong:1-1 and known accessory regions in *Fol*4287 (chr1B; chr2B; chr3; chr6; chr14; chr15)^6^, respectively. Light blue indicates the remaining regions. Ring B: Distribution of transposable elements (TEs) in 50 kb windows. Ring C: Syntenic regions (> 95% identity, 30 kb) between *Fol*4287 and *Focn*Cong:1-1 assemblies (**a**) and between Fo5176 and *Focn*Cong:1-1 assemblies (**b**). Asterisks indicate reverse-complemented scaffolds (SCs) for visual clarity. Ticks on bands represent 1 Mb.

Recently, a chromosome-level genome assembly of the Arabidopsis-infecting *F. oxysporum* isolate Fo5176 was reported^21^. The genomes of *Focn*Cong:1-1 and Fo5176 are very similar, sharing from 93.2% to 94.3% of their total scaffold/contig lengths (> 95% identity, 10 kb). Synteny analysis revealed that (i) SC16 and SC18 of *Focn*Cong:1-1 correspond to chromosome 14 (chr14) of Fo5176, and (ii) SC10 and SC20 to chr16 (Fig. 1b), indicating that these SCs constitute, or contribute to the respective chromosomes. Due to the observations (i) and (ii) above, we refer to the chromosomes carrying these sequences as chr^SC16/SC18^ and chr^SC10/SC20^ in *Focn*Cong:1-1, respectively (see below).

### *Focn*Cong:1-1 has multiple CD chromosomes involved in both vegetative growth and virulence

To identify CD chromosomes of *Focn*Cong:1-1, we generated chromosome-deficient strains by treatment with the mitosis inhibitor benomyl^7, 22^. For the parental strain, we utilized the previously generated strain *Focn*Cong:1-1 Δ*SIX4*, in which *SIX4*, located in SC9, had been replaced with a hygromycin B resistance gene (*hph*) cassette^23^. After benomyl treatment, we obtained six hygromycin B-sensitive mutants (HS1 to HS6; Supplementary Fig. 1). To confirm the loss of dispensable genomic regions, we sequenced the genomes of *Focn*Cong:1-1 Δ*SIX4* and each of the HS mutants. As expected, *Focn*Cong:1-1 Δ*SIX4* maintained SC9 carrying *hph*, but all of the HS mutants had lost SC9 (Fig. 2a, b). We also found that (i) SC3 was absent in HS2, HS3, and HS4, (ii) SC5 and SC8 were lost only in HS6, (iii) chr^SC10/SC20^ was missing in HS3, HS4, HS5, and HS6, and (iv) chr^SC16/SC18^ was lost only in HS4.

**Figure 2.**
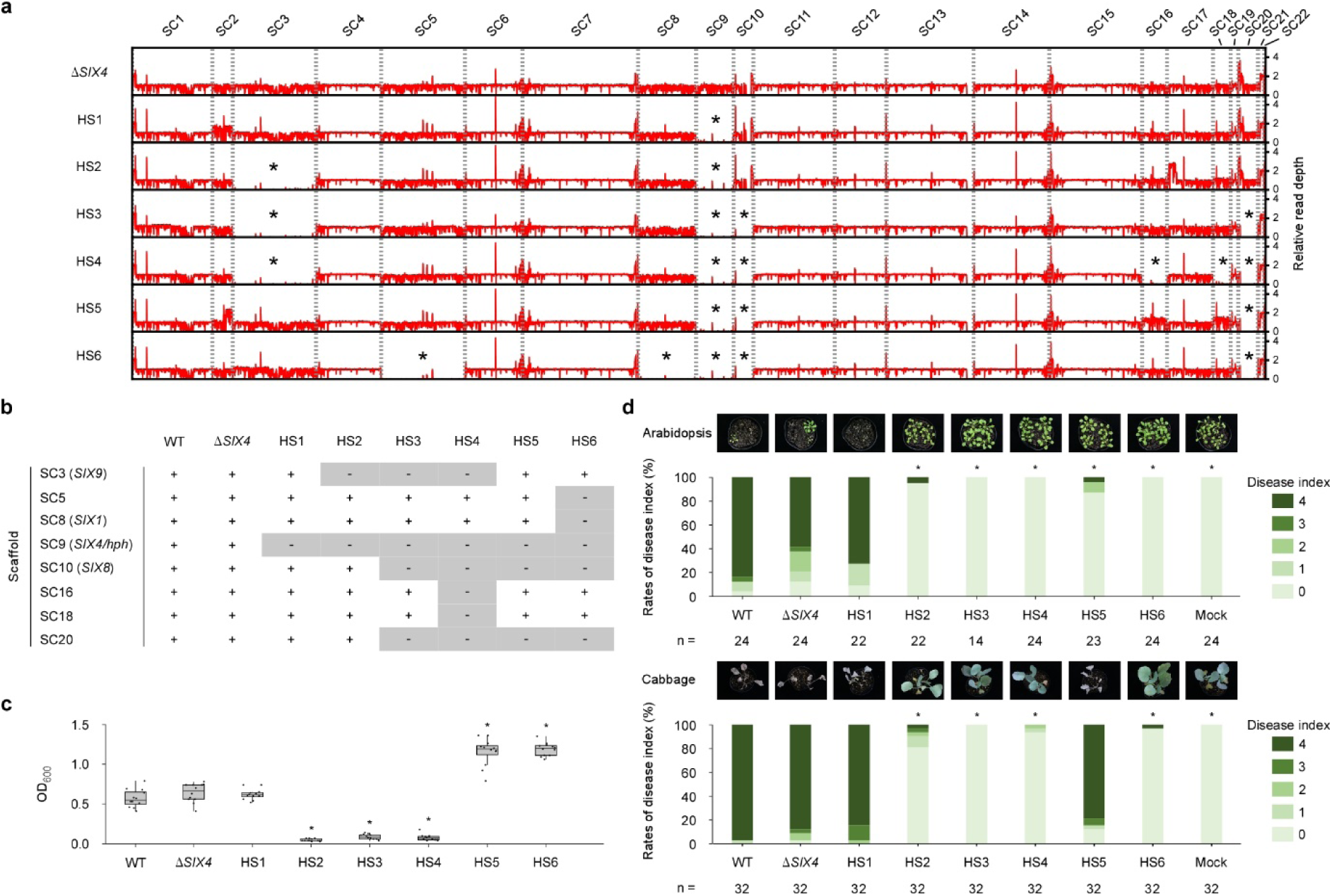
Effects of loss of conditionally dispensable chromosomes on vegetative growth and virulence in *Focn*Cong:1-1. **a,** Relative read mapping depths of *Focn*Cong:1-1 Δ*SIX4* and HSs whole genomes. Asterisks indicate scaffold (SC)-level deletion. **b,** Loss patterns of SC in *Focn*Cong:1-1 HSs. + and – represent maintained- and lost-SCs, respectively. *SIX*s located on particular SCs are shown in parentheses. **c,** Box and whisker plots of conidial formation in *Focn*Cong:1-1 WT, Δ*SIX4* and HSs. OD_600_ of conidial suspensions were measured from six colonies after 17 days of incubation on potato dextrose agar. Results of two independent experiments were combined and a total of twelve biological replicates per isolate are plotted. Asterisks represent significant differences from *Focn*Cong:1-1 Δ*SIX4* (*p < 0.0001, Welch’s t-test). **d,** Virulence of *Focn*Cong:1-1 WT, Δ*SIX4* and HSs to Arabidopsis and cabbage. Disease index was scored as described in Materials and Methods. Results of at least two experiments were combined. ‘n’ denotes the number of plants investigated. Asterisks represent significant differences from *Focn*Cong:1-1 Δ*SIX4* (*p < 0.001, Mann-Whitney U-test). Representative images of Arabidopsis and cabbage at 28 dpi are shown above each bar.

Among the *Focn*Cong:1-1 HS mutants, there was no appreciable difference in colony size, but there was a significant difference in conidial formation (Fig. 2c, and Supplementary Fig. 1c). *Focn*Cong:1-1 HS1 (ΔSC9) showed no difference in conidial formation or virulence on either Arabidopsis or cabbage compared to the parent strain Δ*SIX4*, indicating that SC9 is involved in neither vegetative growth nor virulence (Fig. 2c, d). *Focn*Cong:1-1 mutants without SC3 (HS2, HS3, HS4) had attenuated virulence on Arabidopsis and cabbage, but also had reduced ability to form conidia (Fig. 2c, d), suggesting that SC3 positively regulates conidial formation. To our surprise, loss of chr^SC10/SC20^ in *Focn*Cong:1-1 HS5 and HS6 increased conidial formation (Fig. 2c), but reduced virulence on Arabidopsis (Fig. 2d). Interestingly, *Focn*Cong:1-1 HS6 (ΔSC5/SC8/SC9/chr^SC10/SC20^) lost virulence on both cabbage and Arabidopsis, whereas HS5 (ΔSC9/chr^SC10/SC20^) retained virulence on cabbage, but not on Arabidopsis (Fig. 2d). These data indicate that chr^SC10/SC20^ is required for disease progression on Arabidopsis, but that SC5 and/or SC8 are involved only in causing disease in cabbage. Therefore, SC3, SC5, SC8 and chr^SC10/SC20^ are CD chromosomes affecting disease levels, with SC3 and chr^SC10/SC20^ being also associated with vegetative growth, which is an atypical feature of known CD chromosomes.

### *Focn*Cong:1-1 CD chromosomes are transferable

We investigated whether *Focn*Cong:1-1 CD chromosomes are transferable under laboratory conditions, and what their effect might be on virulence. *Focn*Cong:1-1 HS6 lost multiple virulence-associated CD chromosomes (SC5, SC8, and chr^SC10/SC20^) along with virulence on both Arabidopsis and cabbage (Fig. 2). Strain HS6 could therefore be used to determine the effects of chromosome transfer on virulence. A phleomycin-resistant *Focn*Cong:1-1 HS6 strain (HS6-BLE) was generated by introducing the phleomycin resistance gene (*ble*) (Supplementary Fig. 2a). We co-incubated *Focn*Cong:1-1 HS6-BLE with hygromycin B-resistant *Focn*Cong:1-1 Δ*SIX4* as a donor and selected four colonies (HCT1 to HCT4) that were resistant to both phleomycin and hygromycin B. There was no apparent difference in morphology or colony size (*i.e.* growth rate) in any of the presumptive *Focn*Cong:1-1 recipients (HCT1 to HCT4; Supplementary Fig. 2b). We confirmed chromosome transfer by contour-clamped homogeneous electric field (CHEF) electrophoretic karyotyping as well as PCR (Fig. 3a and Supplementary Fig. 2a). *SIX1* (in SC8) and *hph* (in SC9) were detected in all *Focn*Cong:1-1 recipients, whereas *SIX8* (in SC10) and an SC20 marker (*FocnCong_v011766*) were detected only in HCT1 (Supplementary Fig. 2a). We did not detect the SC5 marker *FocnCong_v016149* in any recipient (Supplementary Fig. 2a). These results indicate that at least SC8, SC9 and chr^SC10/SC20^ are transferable. Conidial formation of the three *Focn*Cong:1-1 recipients HCT2, HCT3 and HCT4, which acquired SC8 and SC9, was comparable to that of HS6-BLE. In contrast, *Focn*Cong:1-1 HCT1, which received chr^SC10/SC20^, SC8 and SC9, produced significantly fewer conidia than HS6-BLE, a phenotype similar to the donor Δ*SIX4* (Fig. 3b). Because SC8 and SC9 are not involved in vegetative growth (Fig. 2c), these results suggest a negative involvement of chr^SC10/SC20^ in vegetative growth. All *Focn*Cong:1-1 recipients (*i.e.* HCT1 to HCT4) that acquired SC8 and SC9 also regained virulence against cabbage (Fig. 3c). Because SC9 is not involved in virulence (Fig.2d), we conclude that SC8 is necessary and sufficient for virulence to cabbage. In the case of Arabidopsis, *Focn*Cong:1-1 HCT1 showed higher virulence than the other recipients (HCT2, HCT3 and HCT4; Fig. 3c). Taken together with the pathology results of the chromosome-deficient *Focn*Cong:1-1 mutants (HS1 to HS6; Fig. 2), it is likely that chr^SC10/SC20^ is required for virulence on Arabidopsis.

**Figure 3.**
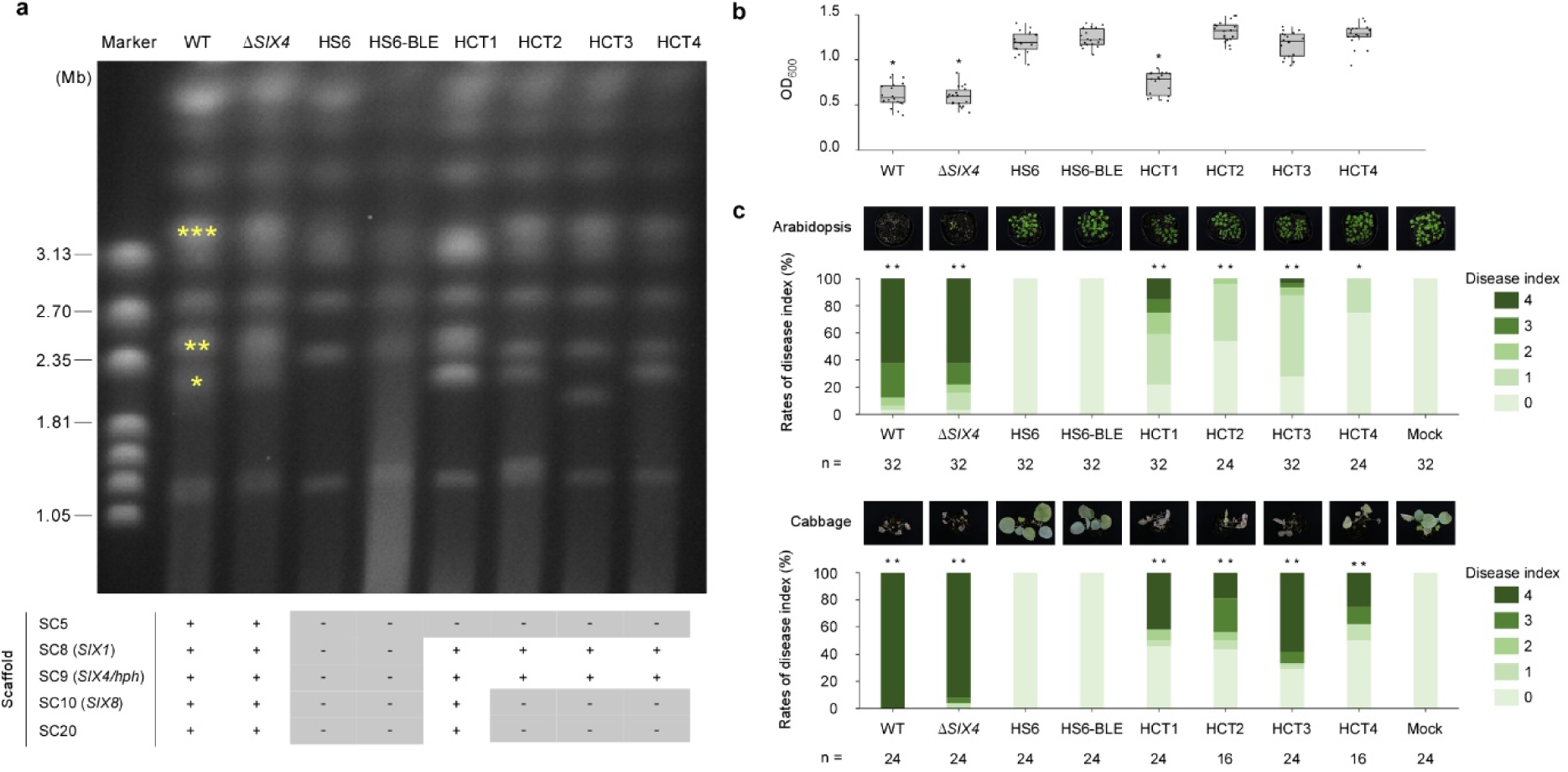
Effects of chromosome transfer on vegetative growth and virulence in *Focn*Cong:1-1 HS6. **a,** Electrophoretic karyotype of *Focn*Cong:1-1 WT, Δ*SIX4,* HS6, HS6-BLE and HCTs. Asterisks indicate chromosomes on which *SIX* genes are located^15^: \**SIX4,* \*\**SIX8*, \*\*\**SIX1*. The table indicates the scaffold (SC) patterns confirmed by PCR as shown in Supplementary Figure 2a. + and – represent maintained- and lost-SCs, respectively. *SIX*s located on SCs are shown in parentheses. **b,** Conidial formation in *Focn*Cong:1-1 WT, Δ*SIX4*, HS6, HS6-BLE and HCTs. OD_600_ of conidial suspension was measured from six colonies after 17 days of incubation on potato dextrose agar. Results of three independent experiments were combined and a total of 18 biological replicates are plotted. Asterisks represent significant differences from *Focn*Cong:1-1 HS6-BLE (*p < 0.0001, Welch’s t-test). **c,** Virulence of *Focn*Cong:1-1 WT, Δ*SIX4*, HS6, HS6-BLE and HCTs on Arabidopsis and cabbage. Disease index was scored as described in Materials and Methods. Results of at least two independent experiments were combined. n denotes the number of plants investigated. Asterisks represent significant difference from *Focn*Cong:1-1 HS6-BLE (**p < 0.001, *p < 0.01 Mann–Whitney U-test). Representative images of Arabidopsis and cabbage at 28 dpi are shown above each bar.

### Chr^SC10/SC20^ is involved in suppression of CYP79B2/CYP79B3-mediated immunity

A CD chromosome from the Arabidopsis-infecting anthracnose fungus *Colletotrichum higginsianum* has been reported to be involved in suppression of plant immunity that is dependent on tryptophan (Trp)-derived secondary metabolites^24^. We investigated whether CD chromosomes of *Focn*Cong:1-1 encode products that also suppress specific immunity. For this experiment, we used the Arabidopsis double mutant *cyp79b2/cyp79b3* that lacks the ability to synthesize Trp-derived secondary metabolites^25^. Among the chromosome-deficient *Focn*Cong:1-1 mutants with attenuated virulence to Arabidopsis Col-0 WT (HS2 to HS6), only *Focn*Cong:1-1 HS5 (ΔSC9/chr^SC10/SC20^) showed the same level of virulence on *cyp79b2/cyp79b3* plants as was observed for its parent strain Δ*SIX4* (Fig. 4a). These results suggest that chr^SC10/SC20^ plays a key role in suppressing Trp-derived secondary metabolite-dependent immunity. *Focn*Cong:1-1 HS6 (ΔSC5/SC8/SC9/chr^SC10/SC20^) was substantially less virulent on *cyp79b2/cyp79b3* plants. This is likely because SC5 or SC8 are involved in virulence other than through suppression of Trp-based immunity. In addition, we found that the *cyp79b2/cyp79b3* double mutant was resistant to SC3-deficient *Focn*Cong:1-1 mutants (HS2 to HS4; Fig 4a), possibly due to some deficiency of vegetative growth in these mutants (Fig. 2c).

**Figure 4.**
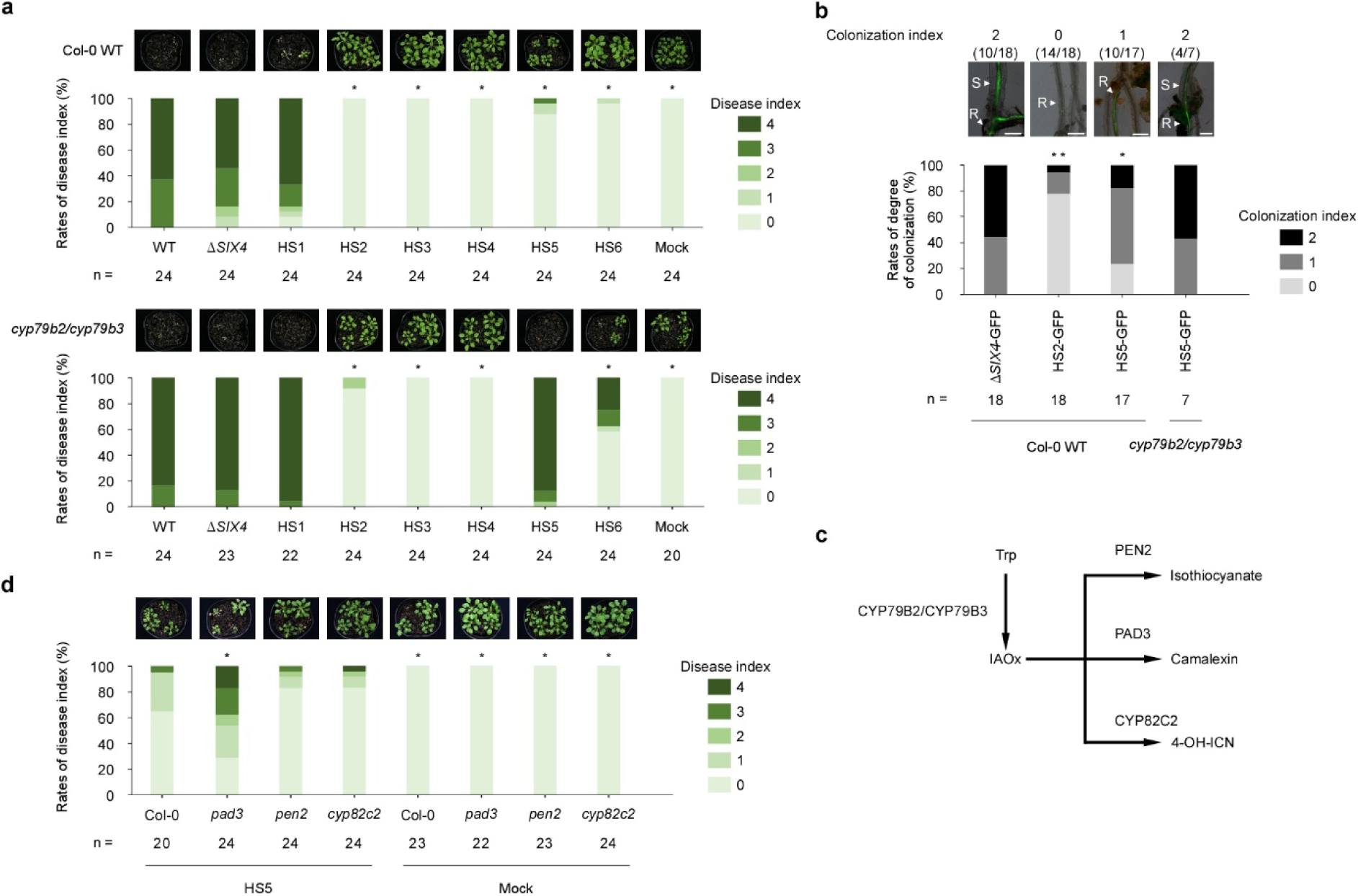
Involvement of chr^SC10/SC20^ in suppression of CYP79B2/CYP79B3-mediated immunity. **a,** Virulence of *Focn*Cong:1-1 WT, Δ*SIX4* and HSs on Arabidopsis Col-0 WT and the *cyp79b2/cyp79b3* double mutant. Disease index was scored as described in Materials and Methods. Results of three independent experiments were combined. n denotes the number of plants investigated. Asterisks represent significant difference from *Focn*Cong:1-1 Δ*SIX4* (*p < 0.001, Mann–Whitney U-test). Representative images of Arabidopsis at 28 dpi are shown above each bar. **b,** Infection phenotypes of *Focn*Cong:1-1 Δ*SIX4*, HS2 and HS5 in Arabidopsis. Colonization index of Arabidopsis Col-0 WT and *cyp79b2/cyp79b3* double mutant inoculated with GFP-labeled *Focn*Cong:1-1 Δ*SIX4* (Δ*SIX4*-GFP), HS2 (HS2-GFP) or HS5 (HS5-GFP) at 12 dpi was scored from 0 to 2: 0, germination or colonization on root surface; 1, colonization in xylem vessels of roots, 2, transition from roots to stems. Results of at least two independent experiments were combined. n denotes the number of plants investigated. Asterisks represent significant differences from *Focn*Cong:1-1 Δ*SIX4*-GFP (**p < 0.001, *p < 0.01, Mann–Whitney U-test). Representative images of root or stem of Arabidopsis at 12 dpi are shown above each bar. Scale bars indicate 200 μm. S, stems; R, roots. **c,** A simple scheme of biosynthesis of tryptophan (Trp)-derived defense compounds in Arabidopsis. IAOx, indole-3-acetaldoxime; 4-OH-ICN, 4-hydroxy-indole carbonyl nitrile. **d,** Virulence of *Focn*Cong:1-1 HS5 on Arabidopsis Col-0 WT, *pad3*, *pen2* and *cyp82c2* mutants. Disease index was scored as described in Materials and Methods. Results of three independent experiments were combined. n denotes the number of plants investigated. Asterisks represent significant difference from Arabidopsis Col-0 WT infected with *Focn*Cong:1-1 HS5 (*p < 0.01, Mann–Whitney U-test). Representative images of Arabidopsis at 28 dpi are shown above each bar.

To investigate which step or steps of infection the CD chromosomes contribute to, a histological analysis was performed using GFP-labeled *Focn*Cong:1-1 strains in Arabidopsis. *Focn*Cong:1-1 Δ*SIX4*-GFP always colonized xylem vessels of roots, often reaching stem elements in Arabidopsis WT, whereas *Focn*Cong:1-1 HS2-GFP lacking SC3 germinated on root surfaces but showed almost no colonization in xylem vessels of roots or stems (Fig. 4b), confirming its deficiency in growth *in planta*. *Focn*Cong:1-1 HS5-GFP (ΔSC9/chr^SC10/SC20^) colonized root xylem vessels, but the frequency of stem colonization was low in Arabidopsis WT. In *cyp79b2/cyp79b3* double mutants, however, *Focn*Cong:1-1 HS5-GFP frequently colonized the stems as was observed for *Focn*Cong:1-1 Δ*SIX4* in WT (Fig. 4b). These results suggest that chr^SC10/SC20^ is implicated in the ability to colonize beyond root xylem vessels into stems, and conversely that CYP79B2/CYP79B3 participate in inhibition of *Focn*Cong:1-1 colonization of stems.

To determine whether CYP79B2/CYP79B3-based antibiotics such as isothiocyanate, camalexin, or 4-hydroxyindole-3-carbonyl nitrile (4-OH-ICN)^26, 27^ are associated with resistance, we conducted bioassays with *Focn*Cong:1-1 HS5 (ΔSC9/chr^SC10/SC20^) on three Arabidopsis mutants (*pen2*, *pad3*, and *cyp82c2*) that are unable to produce isothiocyanate, camalexin, and 4-OH-ICN, respectively^26, 27^ (Fig. 4c). The *pad3* mutant, but not *pen2* or *cyp82c3*, was more susceptible to *Focn*Cong:1-1 HS5 than WT (Fig. 4d), suggesting that camalexin, but not isothiocyanate or 4-OH-ICN, is involved in resistance to *Focn*Cong:1-1. Importantly, camalexin is produced in Arabidopsis, but not in cabbage^28^. Because *Focn*Cong:1-1 HS5 (ΔSC9/chr^SC10/SC20^) had attenuated virulence on Arabidopsis but full virulence on cabbage (Fig. 2d), chr^SC10/SC20^ is likely to contribute to suppression of Arabidopsis-specific immunity, specifically camalexin, to establish infection.

### A pair of effectors are involved in virulence on Arabidopsis

Because chr^SC10/SC20^ is likely to encode effectors that contribute to suppression of Arabidopsis-specific immunity, we searched for genes encoding potential effectors, and found a total of twelve effector candidate genes located on chr^SC10/SC20^ (Supplementary Data 1). Expression profiling revealed that *FocnCong_v001893* (*SIX8*) and *FocnCong_v001894* were highly expressed during infection (Fig 5a and Supplementary Data 2). Interestingly, *SIX8* is adjacent to *FocnCong_v001894,* with an intergenic distance of 1,057 bp on SC10 (Fig. 5b). The intergenic region contains a miniature impala inverted repeat (mimp-IR) sequence, which is related to TE sequences (Fig. 5b and Supplementary Figure 3). mimp-IR is also often located in the upstream regions of *SIX* and other effector candidate genes in *Fol*, *Forc*, and the melon-infecting pathogen *F. oxysporum* f. sp. *melonis*^7, 12, 29^. To determine whether *SIX8* and *FocnCong_v001894* are involved in virulence on Arabidopsis, a genome fragment containing the *SIX8*-*FocnCong_v001894* locus was introduced into *Focn*Cong:1-1 HS5 (ΔSC9/chr^SC10/SC20^) (Supplementary Fig. 4), which restored full virulence to *Focn*Cong:1-1 HS5 (Fig. 5c). In contrast, Arabidopsis WT was resistant to the other *Focn*Cong:1-1 HS5 transformants that contained only *SIX8* or *FocnCong_v001894* (Fig. 5c). It should be noted that virulence of knockout mutants that lack the *SIX8*-*FocnCong_v001894* locus in *Focn*Cong:1-1 was significantly lower than for WT (Supplementary Fig. 5), suggesting that both *SIX8* and *FocnCong_v001894* are necessary for virulence on Arabidopsis. We therefore designated *FocnCong_v001894* as *Pair with SIX Eight1* (*PSE1*).

**Figure 5.**
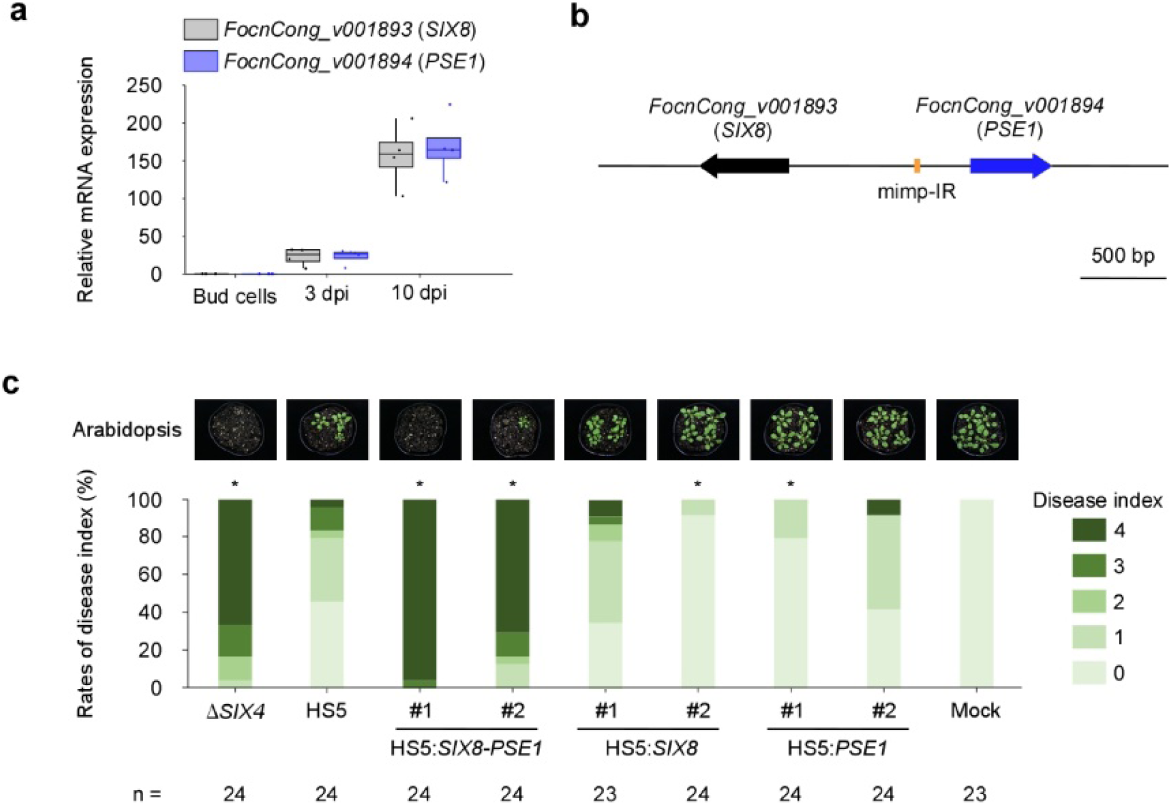
The *SIX8*-*PSE1* locus is involved in virulence of *Focn*Cong:1-1 on Arabidopsis. **a,** Relative transcript levels of *FocnCong_v001893* (*SIX8*) and *FocnCong_v001894* (*PSE1*) in bud cells of *Focn*Cong:1-1 and during infection of Arabidopsis Col-0 WT at 3 and 10 dpi. Expression levels were determined by qRT-PCR and normalized against *Focn*Cong:1-1 *EF1α*. **b,** Schematic representation of the *SIX8*-*PSE1* locus in *Focn*Cong:1-1. mimp-IR, miniature impala-like inverted repeat. **c,** Disease index of Arabidopsis Col-0 WT challenged with *Focn*Cong:1-1 Δ*SIX4,* HS5, HS5 transformants introduced with *SIX8* (HS5:*SIX8*), *PSE1* (HS5:*PSE1*) or both (HS5:*SIX8*-*PSE1*), or water (mock) at 28 dpi was scored as described in Materials and Methods. n denotes the number of plants investigated. Asterisks represent significant difference from *Focn*Cong:1-1 HS5. (*p < 0.001, Mann–Whitney U-test). Representative images of Arabidopsis at 28 dpi are shown above each bar.

### Genetic and functional conservation of the *SIX8* and *PSE1* loci

Next, we investigated whether the *SIX8-PSE1* pair is conserved in Arabidopsis-infecting *F. oxysporum* isolates. Comparative analysis of highly contiguous and available genome assemblies of *F. oxysporum* isolates (Supplementary Table 1)^6, 7, 21, 30–34^ showed that the *SIX8*-*PSE1* locus is completely conserved in Fo5176 and in the stock-infecting pathogen *F. oxysporum* f. sp. *matthiolae* (*Fomt*) PHW726, which can infect Arabidopsis^13, 35^, but not in isolates that cannot infect Arabidopsis (Fig. 6a, b). For example, the banana-infecting pathogen *F. oxysporum* f. sp. *cubense* (*Focb*) tropical race 4 (TR4), which threatens banana production worldwide, has *SIX8* but not *PSE1*. In the other non-Arabidopsis-infecting isolates, except *Fol*4287, neither *SIX8* nor *PSE1* is present. *Fol*4287 has multiple copies of *SIX8* and its homolog *SIX8b*^12, 36^ but *PSE1* is not annotated in the published *Fol*4287 gene annotation^6^. However, we found three loci similar to the *SIX8*-*PSE1* locus in chromosomes 2, 3 and 14 of *Fol*4287. At these loci, adjacent to *SIX8,* there is a *PSE1*-like gene (*PSL1*) differing in the C-terminal 10 amino acids (Fig 6b and Supplementary Figure 6). Furthermore, multiple *SIX8b* loci contain TEs inserted into adjacent *PSE1* sequences. For example, a transposase gene was found in the first intron of the *PSE1* homologs in two loci of chromosome 3 and another locus in chromosome 6 (Fig. 6b and Supplementary Figure 7). Similarly, a presumptive transposase was found immediately upstream of the potential-but-unannotated *PSE1* homolog in another locus in chromosome 6 (Fig. 6b). Thus, TE insertion seems to have disrupted the *PSE1* adjacent to *SIX8b* in *Fol*4287.

**Figure 6.**
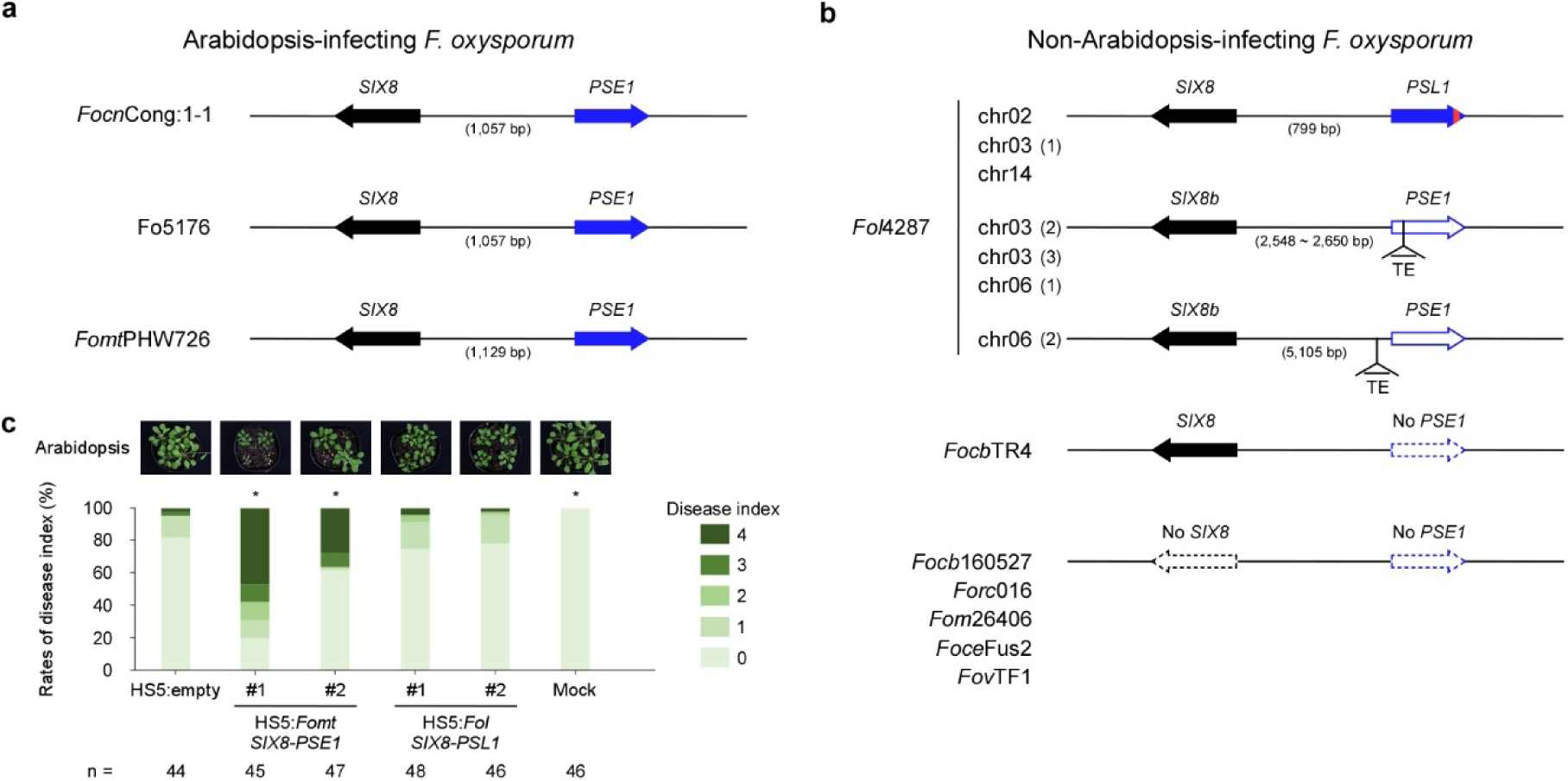
Genetic and functional conservation of a pair of effectors, *SIX8* and *PSE1*, among *F. oxysporum* isolates. Comparison of the *SIX8*-*PSE1* loci in Arabidopsis-infecting (**a**) and non-Arabidopsis-infecting *F. oxysporum* isolates (**b**). A red region in the arrow head of *PSL1* indicates a region of 10 amino acids that differ from *PSE1*. *Focn*Cong:1-1, *F. oxysporum* f. sp. *conglutinans* Cong:1-1; *Fomt*PHW726, *F. oxysporum* f. sp. *matthiolae* PHW726; *Fol*4287, *F. oxysporum* f. sp. *lycopersici* 4287; *Focb*TR4, *F. oxysporum* f. sp. *cubense* tropical race 4; *Focb*160527, *F. oxysporum* f. sp. *cubense* 160527; *Forc*016, *F. oxysporum* f. sp. *radicis-cucumerinum* 016; *Fom*26406, *F. oxysporum* f. sp. *melonis* 26406; *Foce*Fus2, *F. oxysporum* f. sp. *cepae* Fus2; *Fov*TF1, *Fusarium oxysporum* f. sp. *vasinfectum* TF1. **c,** Disease index of Arabidopsis Col-0 WT challenged with *Focn*Cong:1-1 Δ*SIX4*, HS5, HS5 transformants introduced with the *Fomt SIX8*-*PSE1* locus (HS5:*Fomt SIX8*-*PSE1*) or the *Fol SIX8*-*PSL1* locus (HS5:*Fol SIX8*-*PSL1*), or water (mock) at 28 dpi was scored as described in Materials and Methods. Results of six independent experiments were combined. n denotes the number of plants investigated. Asterisks represent significant difference from *Focn*Cong:1-1 HS5. (*p < 0.01, Mann–Whitney U-test). Representative images of Arabidopsis at 28 dpi are shown above each bar.

To evaluate if the *SIX8*-*PSE1* locus in *Fomt*PHW726 and the *SIX8*-*PSL1* locus in *Fol*4287 are able to function similarly in *Focn*Cong:1-1, we cloned these loci and transformed them into the *Focn*Cong:1-1 HS5 mutant (ΔSC9/chr^SC10/SC20^; Supplementary Figure 8). Transformation with the *Fomt SIX8*-*PSE1* locus, but not the *Fol SIX8*-*PSL1* locus, restored full virulence to *Focn*Cong:1-1 HS5 in Arabidopsis Col-0 WT (Fig. 6c). These results suggest that the *SIX8*-*PSE1* locus is functionally distinct from *SIX8*-*PSL1* and is functionally conserved in Arabidopsis-infecting *F. oxysporum* isolates.

## Discussion

Here we report the identification of a CD chromosome in *F. oxysporum* that is required for virulence on Arabidopsis. This CD chromosome encodes a pair of effectors (SIX8 and PSE1) that are involved in suppressing Arabidopsis-specific immunity, and are conserved in the other *F. oxysporum* isolates capable of infecting Arabidopsis. The mode of action potentially involves defense against, or suppression of, the phytoalexin camalexin. We also report that another CD chromosome is required for pathogenicity on cabbage. In addition, certain CD chromosomes are involved in both virulence and vegetative growth.

In plant pathogenic fungi, CD chromosomes associated with virulence are usually not involved in vegetative growth^1, 2^. In this sense, SC3 and chr^SC10/SC20^ in *Focn*Cong:1-1 are atypical CD chromosomes that affect vegetative growth, or at least conidial formation (Fig. 2c). Although the reduced virulence of SC3-deficient *Focn*Cong:1-1 mutants (HS2, HS3 and HS4) on Arabidopsis and cabbage (Fig. 2c, d) may be due to deficiency in the ability to form conidia, or to regulatory step(s) that has multiple unexplored phenotypic effects, we cannot exclude the possibility that yet-unknown effectors located on SC3 are implicated in virulence. Interestingly, SC3 contains a region partly syntenic to chromosome 11, which is a core chromosome of *Fol*4287 (Fig. 1a). This syntenic region may contain dose-effective genes involved in vegetative growth. In contrast to SC3, chr^SC10/SC20^ negatively regulates vegetative growth but positively contributes to virulence on Arabidopsis but not on cabbage (Fig. 2c, d), possibly representing a trade-off between vegetative growth and virulence to a particular host.

*Focn*Cong:1-1 carries multiple CD chromosomes that have distinct virulence functions against specific hosts. For example, the CD chromosome chr^SC10/SC20^-deficient *Focn*Cong:1-1 HS5 is less virulent on Arabidopsis, but is able to develop severe disease on cabbage (Fig. 2d). This result may be explained by the fact that *Focn*Cong:1-1 HS5 maintains the CD chromosome SC8, which harbors a gene, *SIX1*, required for full virulence on cabbage^14^. Consistently, *Focn*Cong:1-1 HS6, which lacks both SC8 and chr^SC10/SC20^, lost pathogenicity on both cabbage and Arabidopsis (Fig. 2d), and introduction of SC8 into HS6 restored virulence on cabbage (Fig. 3c). Thus, we conclude that chr^SC10/SC20^ and SC8 are responsible for host specific virulence on Arabidopsis and cabbage, respectively.

The target of the CD chromosome chr^SC10/SC20^ effector is likely to be *CYP79B2/CYP79B3*-mediated immunity in Arabidopsis, because the loss of chr^SC10/SC20^ attenuated virulence of *Focn*Cong:1-1 HS5 to WT, but not to the *cyp79b2/cyp79b3* double mutant (Fig. 4a). *CYP79B2/CYP79B3* had not previously been implicated in resistance to *F. oxysporum*. For instance, Kidd et al.^37^ reported that susceptibility of *cyp79b2/cyp79b3* to *F. oxysporum* Fo5176 was not different from WT. Consistent with this report, our study shows that virulence of *Focn*Cong:1-1 on *cyp79b2/cyp79b3* is comparable to WT (Fig. 4a). Thus, only the use of CD chromosome-deficient mutants allowed us to uncover the involvement of *CYP79B2/CYP79B3* in resistance to *F. oxysporum*. Furthermore, histological analysis suggests that *CYP79B2/CYP79B3*-mediated immunity may be associated with inhibition of root-stem translocation of *Focn*Cong:1-1 (Fig. 4b). *CYP79B2/CYP79B3* is responsible for synthesis of Trp-derived secondary metabolites, including sulfur-containing compounds that are characteristic of the Brassicaceae^26^. These sulfur-containing antimicrobial compounds differ among Brassicaceae species; for example, camalexin is produced in Arabidopsis, but not in cabbage^28^. Our results suggest that *Focn*Cong:1-1 can overcome the Arabidopsis-specific immunity conferred by *PAD3*, a camalexin synthetic gene (Fig. 4d), when the CD chromosome chr^SC10/SC20^ that encodes the paired effectors *SIX8* and *PSE1* is present. This pair of effectors is highly conserved in Arabidopsis-infecting *F. oxysporum* isolates, but not in other isolates (Fig. 6), thus the presence of a particular CD chromosome that harbors these effector genes would contribute to the determination of host specificity.

We identified *SIX8* and *PSE1* as a gene pair adjacent but encoded on opposite DNA strands (head-to-head orientation) (Fig. 5b and Supplementary Figure 3). Head-to-head orientation of effector genes has been documented for other *SIX* genes in *F. oxysporum*. For instance, in *Fol*, a pair of effector genes *SIX3* (also known as *AVR2*) and *SIX5* are also adjacently located in a head-to-head transcriptional orientation^12, 38, 39^. Both *SIX3* and *SIX5* are required for not only full virulence in a susceptible host, but also disease resistance in tomato lines containing the resistance gene *I-2*^38–40^, and the gene products are thus likely to function as a pair. The close head-to-head orientation may ensure coordinated transcription to produce both proteins at similar levels. Such system would be suitable for maintaining tight stoichiometry of two proteins in a complex. Indeed, SIX5 interacts with SIX3 at plasmodesmata in plant cells, facilitating cell-to-cell movement of SIX3^38, 39^. Unlike the SIX3-SIX5 pair, however, we failed to detect direct interaction between SIX8 and PSE1 in a yeast two-hybrid assay (Supplementary Fig. 9). We cannot exclude the possibility that SIX8 indirectly interacts with PSE1, e.g. via host target(s), or the yeast system may not be suitable for detecting interactions of these proteins. Alternatively, SIX8 and PSE1 may act independently. It is also notable that mutation occurs in only *PSE1*, but not in *SIX8*, in non-Arabidopsis infecting *F. oxysporum* isolates. Perhaps *PSE1*, but not *SIX8*, is recognizable in plants that carry corresponding resistance proteins, thus required to be mutated to avoid detection.

In this work we demonstrate that the host range of *F. oxysporum* depends on CD chromosomes. In this respect, it is interesting that certain isolates, such as *Fol*4287 and *Forc*016, have only a single virulence-associated CD chromosome, whereas *Focn*Cong:1-1 has multiple CD chromosomes, each of which encodes host-specific effectors. Because the *Focn*Cong:1-1 genome is very large (68.8 Mb) compared to most known *F. oxysporum* genomes, such as *Fol*4287 (59.9 Mb)^6^ and *Forc*016 (52.9 Mb)^7^, *Focn*Cong:1-1 is likely to have expanded its host range by acquiring and maintaining additional CD chromosomes. Indeed, Masunaka et al.^41^ have shown that a field isolate of *A. alternata* carrying two putative CD chromosomes has a wide host range. In that case, host-specific toxin genes on different chromosomes determine host range^41^. In the case of *F. oxysporum*, host specificity can be determined, at least in part, by effectors, as seen in this study. Further functional analyses of the *SIX8*-*PSE1* paired effectors and their derivatives will be needed to dissect out the molecular mechanisms underlying effector-based host specificity in *F. oxysporum*.

## Materials and Methods

### Fungal strains and plants

Fungal strains used in this study are listed in Supplementary Table 2^23, 42^. For pre-incubation, all strains were incubated on potato dextrose agar (PDA; Nissui Pharmaceutical Co.) at 28 °C in the dark. For bud cell production, all strains were grown in NO_3_ medium (0.17% yeast nitrogen base without amino acids, 3% sucrose and 1% KNO_3_) at 120 strokes per minute (spm) for 4 days at 28 °C in the dark. For gene expression profiling (Supplementary Data 2), mycelia were harvested after 10 days of incubation on PDA at 28 °C. Bud cells were collected from NO_3_ medium by filtration with a nylon mesh and centrifugation. Hyphae trapped with the nylon mesh were collected. Mycelia from PDA, bud cells, and hyphae were stored at −80 °C until RNA isolation.

Arabidopsis (Col-0 wild type, *pen2*, *pad3*, *cyp82c2* and *cyp79b2/cyp79b3* mutants^26, 27^) and cabbage (cv. Shikidori and cv. Shosyu; Takii Seed) were cultured in pots containing autoclaved Super Mix A (Sakata Seed) and vermiculite (VS kakou). Arabidopsis was grown at 22 °C for 10 h under light and 14 h dark in a growth chamber. Cabbage was grown in a greenhouse.

### Bioassays

For evaluation of disease severity, 14-day-old Arabidopsis and cabbage cv. Shikidori roots were injured with a forceps or a plastic peg and then irrigated with 1 ml of *Focn*Cong:1-1 bud cell suspension (1 × 10^7^ cells/ml). An Arabidopsis disease index was scored at 28 or 29 days post-inoculation (dpi) as: 0, no symptoms; 1, dwarf; 2 yellowing, vein clearing or wilting of one to a few leaves; 3, wilting of a whole plant; 4, dead. A cabbage disease index was also scored at 28 or 29 dpi as: 0, no symptoms; 1, yellowing lower leaves; 2, yellowing lower and upper leaves; 3, whole plant wilting; 4, dead.

For gene expression profiling (Supplementary Data 2), 20- or 21-day-old Arabidopsis and 17-day-old cabbage cv. Shosyu roots were irrigated with 1 ml of bud cell suspension (1 × 10^7^ cells/ml). At 3 dpi and 10 dpi, infected roots were washed with water to remove soil. The roots were stored at −80 °C until RNA isolation.

For observation of colonization of Arabidopsis by *Focn*Cong:1-1, roots of 14-day-old Arabidopsis were cut to approximately 1 cm lengths from the border between roots and stems and soaked in bud cell suspension (1 × 10^7^ cells/ml) for 1 min, then transferred to square plates containing soil. At 12 dpi, roots approximately 5 mm below soil surface were observed by an Olympus BX51 epifluorescence microscopy (Olympus) with excitation of 488 nm for GFP. Images were obtained with an Olympus DP74 digital camera (Olympus) and edited with cellSens (Olympus).

### Fungal growth assays

*Focn*Cong:1-1 strains were grown on PDA for 8 days at 28 °C in the dark from a freezer stock. For measurement of colony diameter, mycelium agar disks were collected from the growing edge of a colony using sterile plastic straws and placed in the center of fresh PDA plates. After 8 days, colony diameter was measured. For quantification of conidial formation, 17-day-old colonies were soaked in 10 ml of water and scraped with a colony spreader. Conidial suspensions were filtrated through a nylon mesh to remove mycelia and conidia were quantified at OD_600_ with a WPA CO 8000 Cell Density Meter (WPA).

### Plasmid construction

Primers used for plasmid construction are listed in Supplementary Table 3. To generate *SIX8*-*PSE1* locus complementation vectors, the *Focn*Cong:1-1 *SIX8*-*PSE1*, *Fomt SIX8*-*PSE1* and *Fol SIX8*-*PSL1* loci were amplified from genomic DNAs of *Focn*Cong:1-1, *Fomt*MAFF240332 and *Fol*4287, respectively, and cloned into pCR™8/GW/TOPO® vector using a pCR™8/GW/TOPO® TA Cloning® Kit (Invitrogen) as described by the manufacturer. To introduce these loci into *Focn*Cong:1-1 HS5, the complementation vector containing each locus was co-transformed with pCSN43 containing an *hph* cassette^43^. For transformation vectors of *SIX8* or *PSE1*, an *hph* cassette was amplified from pCSN43^43^ and assembled with *SIX8* or *PSE1*, which was amplified from the *Focn*Cong:1-1 *SIX8*-*PSE1* locus complementation vector as a template, using NEBuilder HiFi DNA Assembly Master Mix (New England Biolabs) as recommended by the manufacturer. The assembled fragments were cloned into pCR™8/GW/TOPO®.

To generate the *Focn*Cong:1-1 *SIX8*-*PSE1* locus disruption vector, the flanking regions of *SIX8* and *PSE1* were amplified from the *Focn*Cong:1-1 *SIX8*-*PSE1* locus complementation vector as a template and assembled with an *hph* cassette using NEBuilder HiFi DNA Assembly Master Mix (New England Biolabs). The assembled fragment was cloned into pCR™8/GW/TOPO®.

Constructs for yeast two-hybrid assays were generated from cDNAs of *SIX8* and *PSE1* without signal peptide sequences or stop codon by amplification from cDNA generated from mRNA isolated from *Focn*Cong:1-1-infected Arabidopsis. Amplicons were inserted into pENTR™/D-TOPO® (Invitrogen), and then into yeast expression vectors pDEST-DB and pDEST-AD^44^ using Gateway™ LR Clonase™ II Enzyme Mix (Invitrogen) as described by the manufacturer.

### Protoplast formation and transformation

*Focn*Cong:1-1 bud cells (4 × 10^8^) were incubated in 80 ml of potato dextrose broth (Difco) at 80 spm for 16 h at 28 °C. Germinated mycelia were collected by centrifugation (1,800*g*, 10 min) and washed with 1.2 M MgSO_4_. Mycelial cell walls were digested with 25 ml 2% (w/v) Driselase (Sigma) and 2% (w/v) Lysing Enzymes (Sigma) in 1.2 M MgSO_4_, and maintained at 80 spm for 3 h at 28 °C. Protoplasts were collected by filtration with a nylon mesh and centrifugation (1,500*g* 10 min), and rinsed twice with 0.7 M NaCl. The protoplasts were resuspended in STC (1.2 M sorbitol, 50 mM CaCl_2_, 10 mM Tris–HCl pH 7.5) and adjusted to 1 × 10^8^ cells/ml. For polyethylene glycol transformation, 30 μg plasmid DNA was added to 150 μl of the protoplast suspension as previously described^44^. Transformants were selected and maintained on PDA containing hygromycin B (100 μg/ml) or G418 (200 μg/ml) and verified by PCR using primers listed in Supplementary Table 3. Plasmid DNAs used for transformation are shown in Supplementary Table 4^43, 45, 46^.

### Genome sequencing and assembly

For PacBio sequencing, genomic DNA of *Focn*Cong:1-1 was isolated using CTAB and 100/G genomic tips (QIAGEN) as described in the 1000 Fungal genomes project (http://1000.fungalgenomes.org). The genome was sequenced on five PacBio RSII cells and assembled by the Hierarchical Genome Assembly Process (HGAP) v4 within SMRT Link (v5.1.0). Default values were kept and the expected genome size was set to 70 Mb.

For optical mapping, genomic DNA was isolated using a Blood and Cell Culture DNA Isolation Kit (Bionano Genomics) as described by the manufacturer. Genomic DNA was labelled with an NLRS Labeling Kit (Bionano Genomics) with *Bsp*QI and *Bbv*CI as described by the manufacturer. The labeled DNA was scanned using a Bionano Irys platform. Bionano maps from two enzymes (*Bsp*QI and *Bbv*CI) (Bionano Solve v3.2) were merged with PacBio sequence assemblies to produce long hybrid scaffolds. Completeness of gene space within the assembly was assessed through the presence of conserved single-copy genes using BUSCO version 3.0.2^47, 48^. Analysis with the Sordariomyceta data set (3,725 genes) indicated the presence of 3,690 genes (99.1%) in the assembly (Table 1). Whole-genome alignments were performed with nucmer (with –maxmatch) in MUMmer 3.23^49^.

For genome sequencing of *Focn*Cong:1-1 Δ*SIX4* and HSs, genomic DNA was isolated using DNeasy Plant Mini Kits (QIAGEN). Illumina NovaSeq 6000 or HiSeq 2500 paired-end sequencing was used for *Focn*Cong:1-1 Δ*SIX4* and HSs, except for HS3, using a library with a mean insert size of 550 bp. Illumina NextSeq 500 single-end sequencing was used for *Focn*Cong:1-1 HS3, from library preparation with a mean insert size of 350 bp. The Illumina sequence library was quality-filtered using the FASTX Toolkit 0.0.13.2 (Hannonlab) with parameters -q20 and -p50. Reads containing “N” were discarded. Quality-filtered libraries were aligned with the *Focn*Cong:1-1 genome using CLC Genomic Workbench 20 using default settings.

### RNA extraction, cDNA synthesis, and qRT-PCR

Total RNAs were extracted using RNeasy Plant Mini Kit (QIAGEN). One μg total RNA was used to generate cDNA in a 20 μl volume reaction according to the Invitrogen Superscript III Reverse Transcriptase protocol. cDNA was diluted 1:5, and 1 μl was used for a 10 μl qPCR reaction with 5 μl THUNDERBIRD SYBR Green mix (Toyobo) on an Mx3000P qPCR System (Agilent) using the following program: (1) 95 °C, 1 min, (2) [95 °C, 15 s, then 53 °C, 30 s, then 72 °C, 1 min] × 40, (3) 95 °C, 1 min for *SIX8* and *PSE1*, or (1) 95 °C, 3 min; (2) [95 °C, 30 s, then 55 °C, 30 s, then 72 °C, 30 s] × 40, (3) 95 °C, 1 min for *Focn*Cong:1-1 *EF1α,* followed by a temperature gradient from 55 to 95 °C. Standard curves were generated using serial dilutions of cDNAs from Arabidopsis infected with *Focn*Cong:1-1 at 10 dpi for *SIX8* and *PSE1* and cDNAs from bud cells for *Focn*Cong:1-1 *EF1α*. *Focn*Cong:1-1 *EF1α* was used as a reference gene. Primers used for qPCR are listed in Supplementary Table 3^50^.

### RNA sequencing

Using the extracted RNA, strand-specific shotgun type of RNA library was prepared using the Breath Adapter Directional sequencing protocol^51^. Briefly, mRNA was extracted and fragmented using magnesium ions at elevated temperature. The polyA tails of mRNA was primed by an adapter-containing oligonucleotide for cDNA synthesis. 5’ adapter addition is performed by breath capture technology to generate strand-specific libraries. The final PCR enrichment was performed using oligonucleotides containing the full adapter sequence with different indexes. After cleanup and size selection, concentration of library was measured by microplate photometer Infinite® 200 PRO (TECAN) to pool libraries for Illumina sequencing systems. The libraries were sequenced on an Illumina NextSeq 500 platform. The Illumina sequence library was quality-filtered and aligned as above. Transcription levels for each transcript were calculated as TPM (transcripts per million).

### Gene prediction and annotation

RNA sequencing data from *Focn*Cong:1-1 was aligned with the *Focn*Cong:1-1 genome using HISAT2 v.2.1.0^52^ and used to guide gene model prediction using the BRAKER1 v1.9 pipeline^53^. BRAKER1 was run with the repeat-softmasked genome created by RepeatMasker v.4.0.7 (with -engine ncbi -species “ascomycota” –xsmall; http://www.repeatmasker.org/), using the fungal and softmasking options. Gene-coding sequences were annotated through BLASTp (E-value cutoff = 1E-6) searches against the July 2018 release of the SWISS-PROT database^54^. Putative secreted proteins were identified through prediction of signal peptides using SignalP v.5.0^55^ and removal of sequences with TMHMM v.2.0^56^-predicted transmembrane domains. For effector prediction, putative secreted proteins were screened for proteins with an effector-like structure using EffectorP 1.0 and/or 2.0^17, 18^. In addition, BLASTp analyses (E-value cutoff = 1E-6) were performed for the fourteen *SIX* genes (*SIX1-SIX14*) and the four genes (*FOA1-FOA4*) known to be effectors in Arabidopsis-infecting *F. oxysporum*^12, 16^.

### Analysis of repeat elements

Repeat element prediction was performed using the genome sequences of eight *F. oxysporum* strains in the NCBI database that had contig N50 values greater than 1 Mb (last accessed on November 24, 2019) as described in Gan et al.^57^. Code used for this analysis are available at: https://github.com/pamgan/colletotrichum_genome. The details of genome sequences used for this analysis are shown in Supplementary Table 5^7, 31–33, 58, 59^. Briefly, repeat sequences were predicted using RECON and RepeatScout via RepeatModeler open-1.0.11 (http://www.repeatmasker.org), TransposonPSI (http://transposonpsi.sourceforge.net/), LTR_retriever^60^, and LTRPred^61^ (https://github.com/HajkD/LTRpred). Sequences that were longer than 400 bp from TransposonPSI, LTR_retriever, and LTRPred were combined and used as queries for BLASTx against RepBase^62^ peptide sequences bundled in RepeatMasker open-4.0.9-p2 (http://www.repeatmasker.org). Lastly, these sequences were used as queries for BLASTn against each fungal genome. Only sequences with more than five hits (BLASTn E-value cutoff = 1E-15) and/or with a hit to a RepBase peptide (BLASTx E-value cutoff = 1E-5) were retained for further analysis. Sequences from all sources were combined using VSEARCH v2.14.0^63^, using 80% identity as the cutoff threshold. Consensus sequences were classified using RepeatClassifier (from RepeatModeler open-1.0.11). Known *Fusarium* repeat sequences registered in Dfam_Consensus-20181026 and RepBase-20181026 were extracted, except for those that were annotated as artefacts, simple repeats, or low complexity sequences. The custom repeat library was created by combining the consensus sequences and known *Fusarium* repeat sequences, and used as input for RepeatMasker open-4.0.9-p2. The “one code to find them all”^64^ was used to reconstruct repeat elements.

### Chromosome loss and transfer

A chromosome loss experiment was performed according to VanEtten et al.^22^. *Focn*Cong:1-1 Δ*SIX4* was incubated in M100 medium (1% glucose, 0.3% KNO_3_, 2% agar, 6.25% salt solution) with benomyl (1.56, 3.13 or 6.25 μg/ml) at 120 spm for 4 days at 28 °C. The salt solution consisted of 0.4% KH_2_PO_4_, 0.4% Na_2_SO_4_, 0.8% KCl, 0.2% MgSO_4_·2H_2_O, 0.1% CaC1_2_, and 0.8% trace elements (0.006% H_3_BO_3_, 0.014% MnCl_2_·4H_2_O, 0.0844% ZnSO_4_·7H_2_O, 0.004% NaMoO_4_·2H_2_O, 0.006% FeCl_3_, 0.04% CuSO_4_·5H_2_O). Hyphae were removed with a nylon mesh, and bud cells were collected by centrifugation at 1,630*g* for 10 min. Supernatant was discarded and the remnant with bud cells was spread on M100 plates containing 0.04% Triton X-100 (Wako), and the inoculated plate was overlaid with an autoclaved filter paper. Plates were incubated at 28 °C for 1 to 3 days, then the filter paper was transferred onto M100 medium containing hygromycin B (100 μg/ml) and incubated at 25 °C overnight. Hygromycin B-sensitive isolates were selected by comparing the plates, and then chromosome loss patterns were verified by PCR (Supplementary Figure 1) using primers listed in Supplementary Table 3^65, 66^.

Chromosome transfer experiments were performed according to van der Does and Rep^67^. A zeocin-resistant *Focn*Cong:1-1 HS6 (HS6-BLE) strain was generated by *Agrobacterium*-mediated transformation as previously reported^68^ with *Agrobacterium tumefaciens* EHA105 harboring pRW1p^69^. *Focn*Cong:1-1 Δ*SIX4* and HS6-BLE were co-incubated on PDA at 25 °C. Conidia were harvested from 7-day-old colonies, and conidial suspensions were spread on PDA containing hygromycin B (100 μg/ml) and phleomycin (100 μg/ml). Double drug-resistant colonies were selected, and then chromosome patterns were verified by PCR (Supplementary Figure 2) using primers listed in Supplementary Table 3^65, 66^.

### Contour-clamped homogeneous electric field (CHEF) gel electrophoresis

CHEF gel plugs were made by resuspending protoplasts in STE (1 M sorbitol, 25 mM Tris-HCl pH 7.5, 50 mM EDTA). Protoplast concentration was adjusted to 4 × 10^8^ cells/ml and added to the same amount of 1.2% low melting agarose gel (Bio-Rad) solution. Protoplast suspensions (2 × 10^8^ cells/ml) containing 0.6% low melting agarose gel were added to 50-well dispensable mold plates (Bio-Rad). Plugs were immersed in 10 ml of NDS (1% N-lauroyl sarcosinate sodium salt solution, 0.01 M Tris-HCl, 0.5 M EDTA) and incubated at 65 spm for 14 to 20 h at 37 °C. NDS was replaced with 0.05 M EDTA three times every 30 min. Plugs in 0.05 M EDTA were stored at 4 °C until use.

CHEF gel electrophoresis was done according to Inami et al.^70^. Briefly, chromosomes were separated on 1% SeaKem® Gold Agarose (Lonza) in 0.5×TBE buffer at 4 to 7 °C for 260 h using a CHEF Mapper System (Bio-Rad). Switching time was 1,200 to 4,800 s at 1.5 V/cm with an included angle of 120°. The running buffer was exchanged every two or three days. Chromosomes of *Hansenula wingei*, *Saccharomyces cerevisiae* and *Schizosaccharomyces pombe* (Bio-Rad) were used as DNA size markers. Gels were stained with 3×GelGreen (Biotium) to visualize chromosomes.

### Yeast two-hybrid assays

For yeast two-hybrid assays, bait (pDEST-DB; DB) and prey vectors (pDEST-AD; AD) containing cDNA of *SIX8*, *PSE1* or empty vector controls were transformed into *S. cerevisiae* Y8930 and Y8800, respectively, with a slight modification of the method described by Lopez and Mukhtar et al.^71^. Transformants carrying DB and AD were selected with synthetic dropout (SD) media (0.67% yeast nitrogen base, 0.5% glucose, 0.01% adenine hemisulfate salt) supplemented with -Leu DO supplement (Clontech) (SD-Leu) and -Trp DO supplement (Clontech) (SD-Trp), respectively. Yeast transformants were mated in yeast extract peptone dextrose growth broth (1% yeast extract, 2% peptone, 2% glucose, 0.01% adenine hemisulfate salt) at 150 spm for 24 h at 28 °C. Diploid cells were selected with SD supplemented with -Leu/-Trp DO supplement (Clontech) (SD-Leu-Trp), and spotted on SD supplemented with -Leu/-Trp/-His DO supplement (Clontech) (SD-Leu-Trp-His) and SD-Leu-Trp with 1 mM 3-amino-1,2,4-triazole. Yeast colonies were observed after 72 h incubation.

### Statistical analysis

All statistical analyses were performed in EZR^72^.

## Data Availability

The Whole Genome Shotgun project of *Focn*Cong:1-1 has been deposited at DDBJ/ENA/GenBank under the accession RSAI00000000 (BioProject number PRJNA506492 and BioSample number SAMN10461798). The version described in this paper is version RSAI01000000. RNA sequencing data from culture medium and plant infections have been deposited in NCBI’s Gene Expression Omnibus (GEO) and are accessible through GEO Series accession number GSE157823.

## Acknowledgments

We thank Prof. Yoshitaka Takano (*pen2* and *pad3*), Dr. Elizabeth S. Sattely (*cyp82c2*), and Dr. Kei Hiruma (*cyp79b2/cyp79b3*) for providing seeds. We also thank Dr. M. Shahid Mukhtar for providing pDEST-DB and pDEST-AD vectors. This work was supported by JSPS KAKENHI 19H00939 (S.A. and T.A.), 20H02995 (S.A.), 17K07679 (S.A.), 19K21154 (Y.A.), and 17H06172 (K.S.); JST PRESTO Grant Number JPMJPR16O1 (S.A.); the Institute for Fermentation, Osaka (Y.A.); research fellowship from the Japan Society for the Promotion of Science (Y.A.).

## Author Contributions

Y.A., S.A., P.G., A.T., I.Y. and A.S. conducted experiments. Y.A., S.A., K.K., P.M.H., M.R., K.S. and T.A. conceived and supervised the study. Y.A., S.A., K.S. and T.A. wrote the manuscript. All authors reviewed and approved the manuscript.

---

**Supplementary Figure 1.**
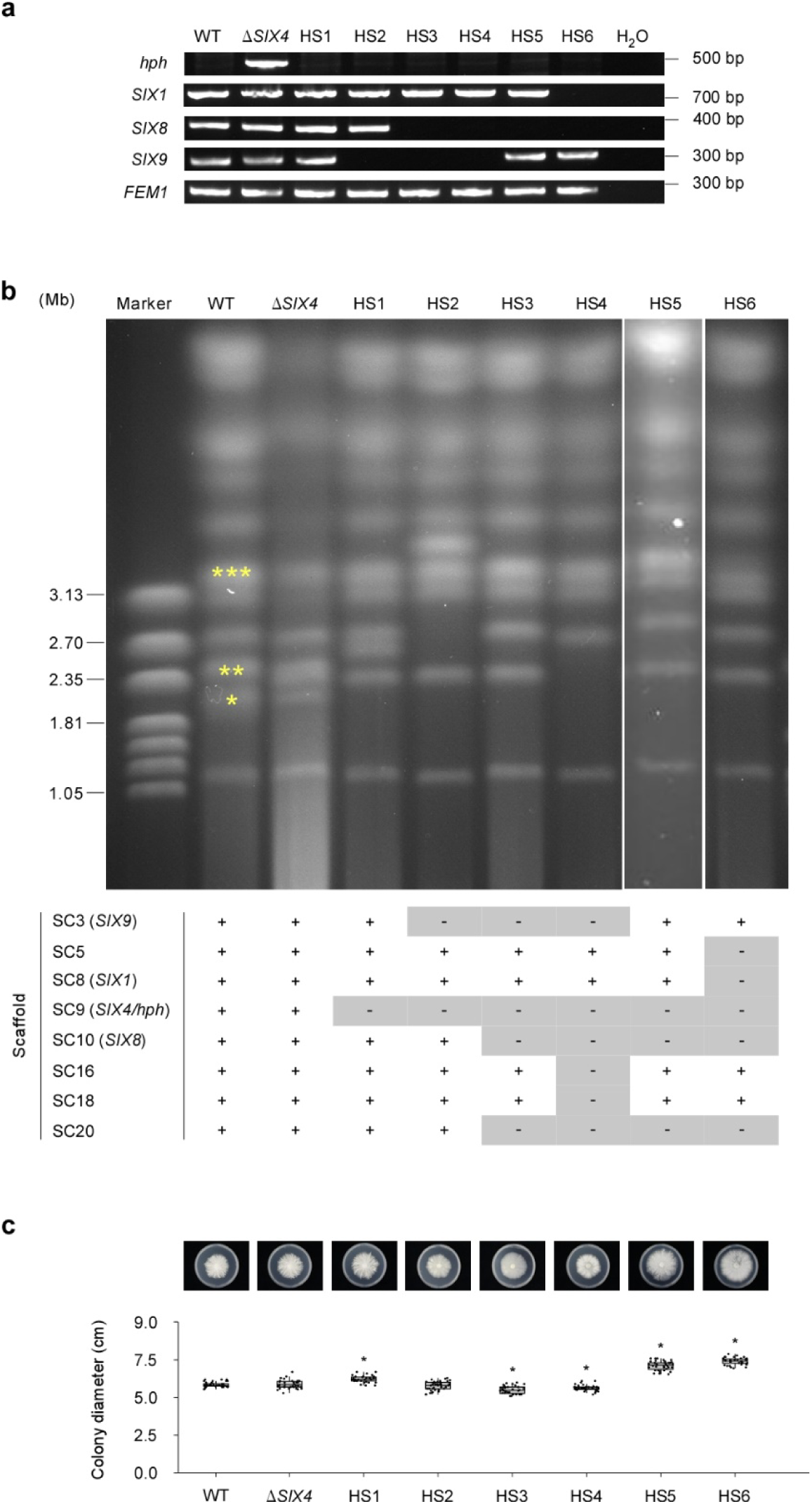
Effects of loss of dispensable chromosomes on radial growth in *Focn*Cong:1-1. **a,** Detection patterns of *SIX* genes in hygromycin B-sensitive *Focn*Cong:1-1 mutants (HS1 to HS6). Hygromycin B resistance gene (*hph*), *SIX1*, *SIX8* and *SIX9* genes were detected by PCR. *FEM1* amplification was used as a control. DNAs of *Focn*Cong:1-1 WT, Δ*SIX4* and HSs were used as a template. **b,** Electrophoretic karyotype of *Focn*Cong:1-1 WT, Δ*SIX4* and HSs. Asterisks indicate chromosomes on which *SIX* genes are located as follows: \**SIX4*, \*\**SIX8*, \*\*\**SIX1*. SC loss patterns estimated by genome sequencing described in Fig. 2a are shown as a table. + and – represent maintained- and lost-SCs, respectively. *SIX*s located on SCs are shown in parentheses. **c,** Colony formation of *Focn*Cong:1-1 WT, Δ*SIX4* and HSs on potato dextrose agar. Results of six independent experiments were combined and a total of 35 or 36 biological replicates are plotted. Asterisks represent significant difference from Δ*SIX4* (*p < 0.0001, Welch’s t-test). Representative images of colonies after 8 days of incubation are shown above each box plot.

**Supplementary Figure 2.**
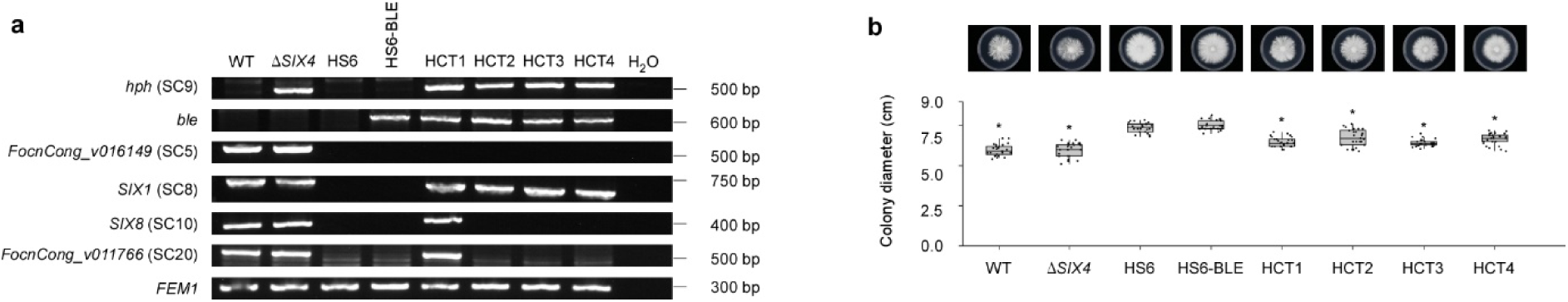
Effects of chromosome transfer on radial growth in *Focn*Cong:1-1 HS6. **a,** Detection patterns of genes located on particular scaffolds (SCs) in horizontally chromosome-transferred recipient colonies (HCT1 to HCT4). *hph*, zeocin resistance gene (*ble*), *FocnCong_v016149*, *SIX1*, *SIX8* and *FocnCong_v011766* were detected by PCR. *FocnCong_v016149*, *SIX1*, *SIX8* and *FocnCong_v011766* are located on SC5, SC8, SC10 and SC20, respectively. *FEM1* was amplified as a control. DNAs of *Focn*Cong:1-1 WT, Δ*SIX4,* HS6, HS6-BLE and HCTs were used as templates. **b,** Colony formation of *Focn*Cong:1-1 WT, Δ*SIX4,* HS6, HS6-BLE and HCTs on potato dextrose agar. Results of four independent experiments were combined and a total of 25 biological replicates are plotted. Asterisks represent significant difference from HS6-BLE (*p < 0.0001, Welch’s t-test). Representative images of colonies after 8 days of incubation are shown above each box plot.

**Supplementary Figure 3.**
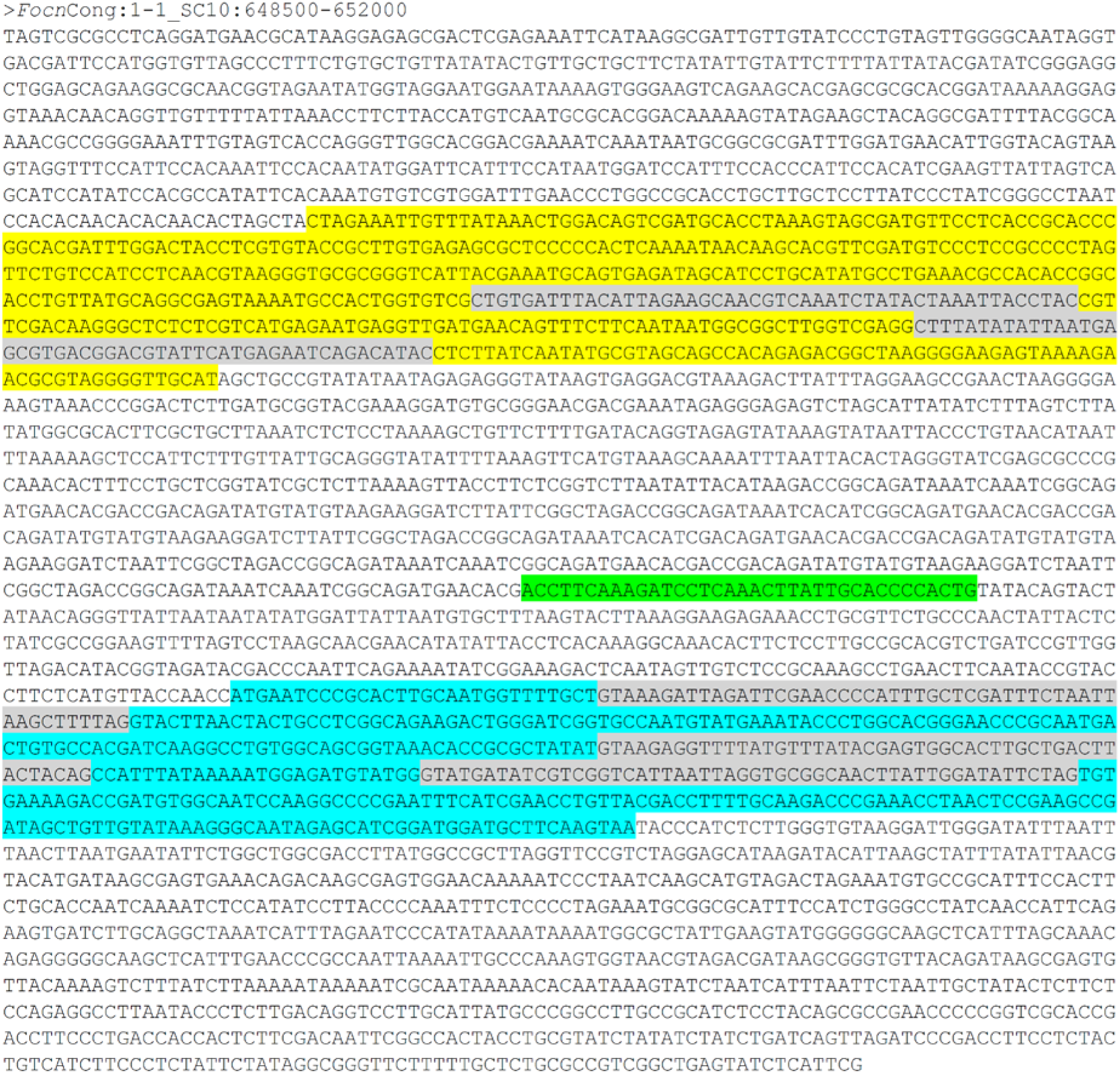
Nucleotide sequence of the *SIX8*-*PSE1* locus of *Focn*Cong:1-1. Yellow and vivid blue highlights indicate the *SIX8* and *PSE1* coding sequences (CDS) in opposite transcriptional orientation. Intron sequences are highlighted in gray. Green highlight represents the miniature impala-like inverted repeat (mimp-IR) sequence.

**Supplementary Figure 4.**
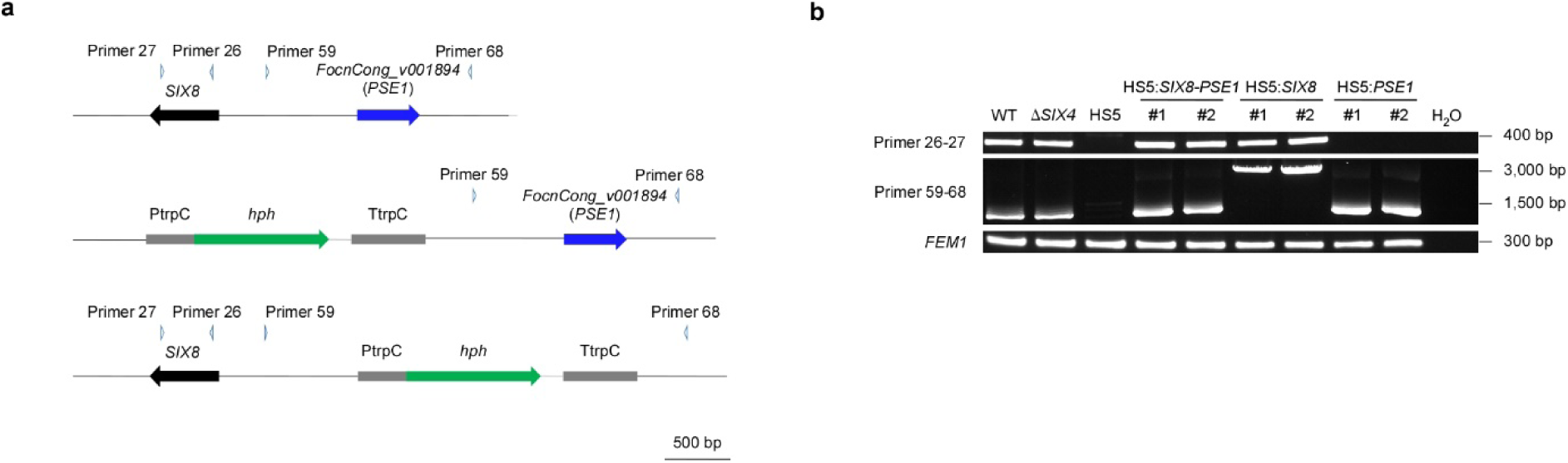
Introduction of the *SIX8*-*PSE1* locus, *SIX8* or *PSE1* into *Focn*Cong:1-1 HS5. **a,** Schematic representation of the *SIX8*-*PSE1* locus (upper) and the loci in which *SIX8* (middle) or *PSE1* (lower) are replaced by an *hph* cassette. Arrowheads indicate the primer locations for PCR verification. **b,** Confirmation of *Focn*Cong:1-1 HS5 transformants introduced with the *SIX8*-*PSE1* locus (HS5:*SIX8*-*PSE1*), *SIX8* (HS5:*SIX8*) or *PSE1* (HS5:*PSE1*) by PCR. *FEM1* was amplified as a control. DNAs of *Focn*Cong:1-1 WT, Δ*SIX4*, two independent HS5:*SIX8*-*PSE1*strains (#1, #2), HS5:*SIX8* strains (#1, #2) and HS5:*PSE1* strains (#1, #2) were used as templates.

**Supplementary Figure 5.**
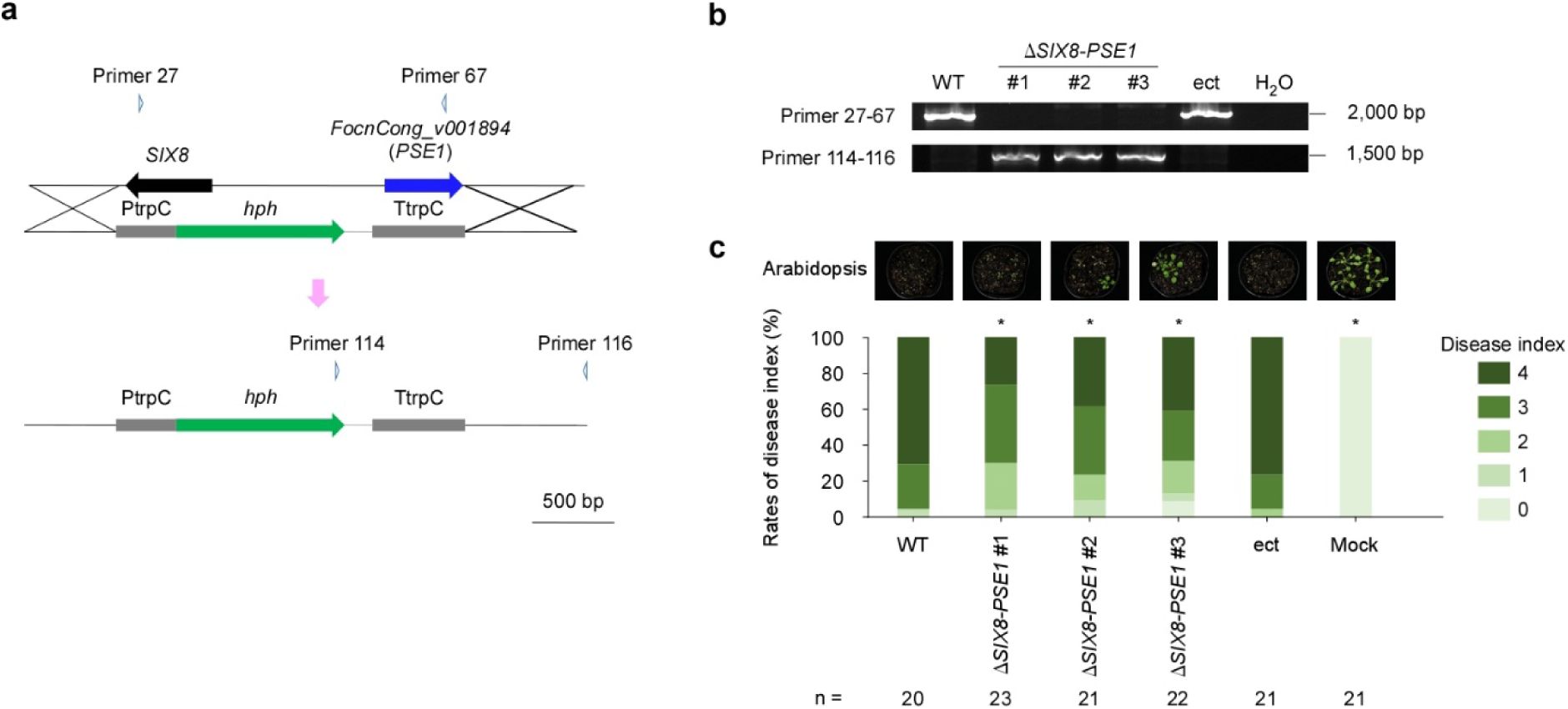
Effects of deletion of the *SIX8-PSE1* locus on virulence of *Focn*Cong:1-1 on Arabidopsis. **a,** Schematic representation of a knockout of the *SIX8-PSE1* locus via replacement with an *hph* cassette. Arrowheads indicate the primer locations for knockout verification. **b,** Confirmation of the *SIX8-PSE1* locus knockout by PCR. DNAs of *Focn*Cong:1-1 WT, three independent *SIX8*-*PSE1* knockout mutants (Δ*SIX8-PSE1* #1-3) and an ectopic transformant (ect) were used as templates. **c,** Disease index of Arabidopsis Col-0 WT challenged with *Focn*Cong:1-1 WT, Δ*SIX8*-*PSE1*, ect or water (mock) at 28 dpi was scored as described in Materials and Methods. Results of three independent experiments were combined. n denotes the number of plants investigated. Asterisks represent significant difference from WT (*p < 0.05, Mann–Whitney U-test). Representative images of Arabidopsis at 28 dpi are shown above each bar graph.

**Supplementary Figure 6.**
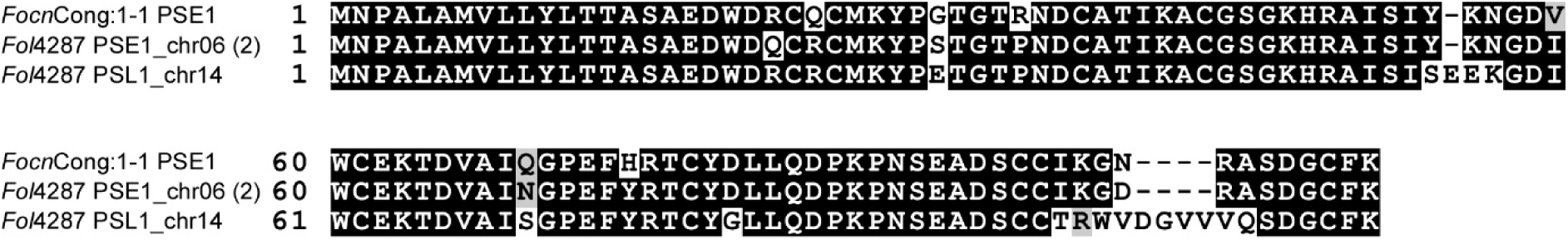
Alignment of the predicted amino acid sequences of PSE1 alleles. Identical sequences are indicated in white on black, and similar amino acids in hydrophobic or hydrophilic features are highlighted in black on gray. Dashes indicate gaps introduced to maximize alignment. Multiple alignments of the amino acid sequences were made using the Clustal Omega method (https://www.ebi.ac.uk/Tools/msa/clustalo/).

**Supplementary Figure 7.**
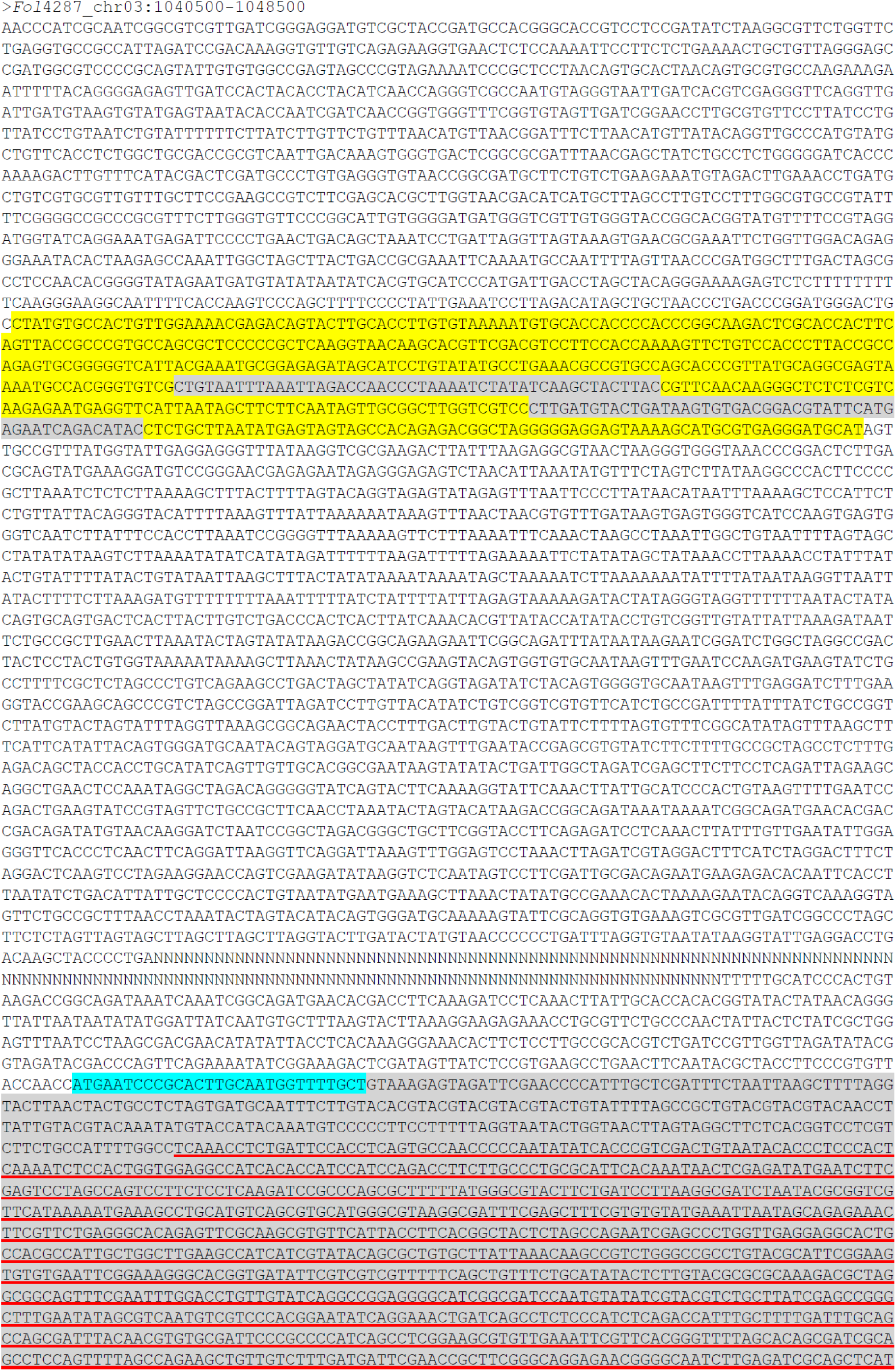
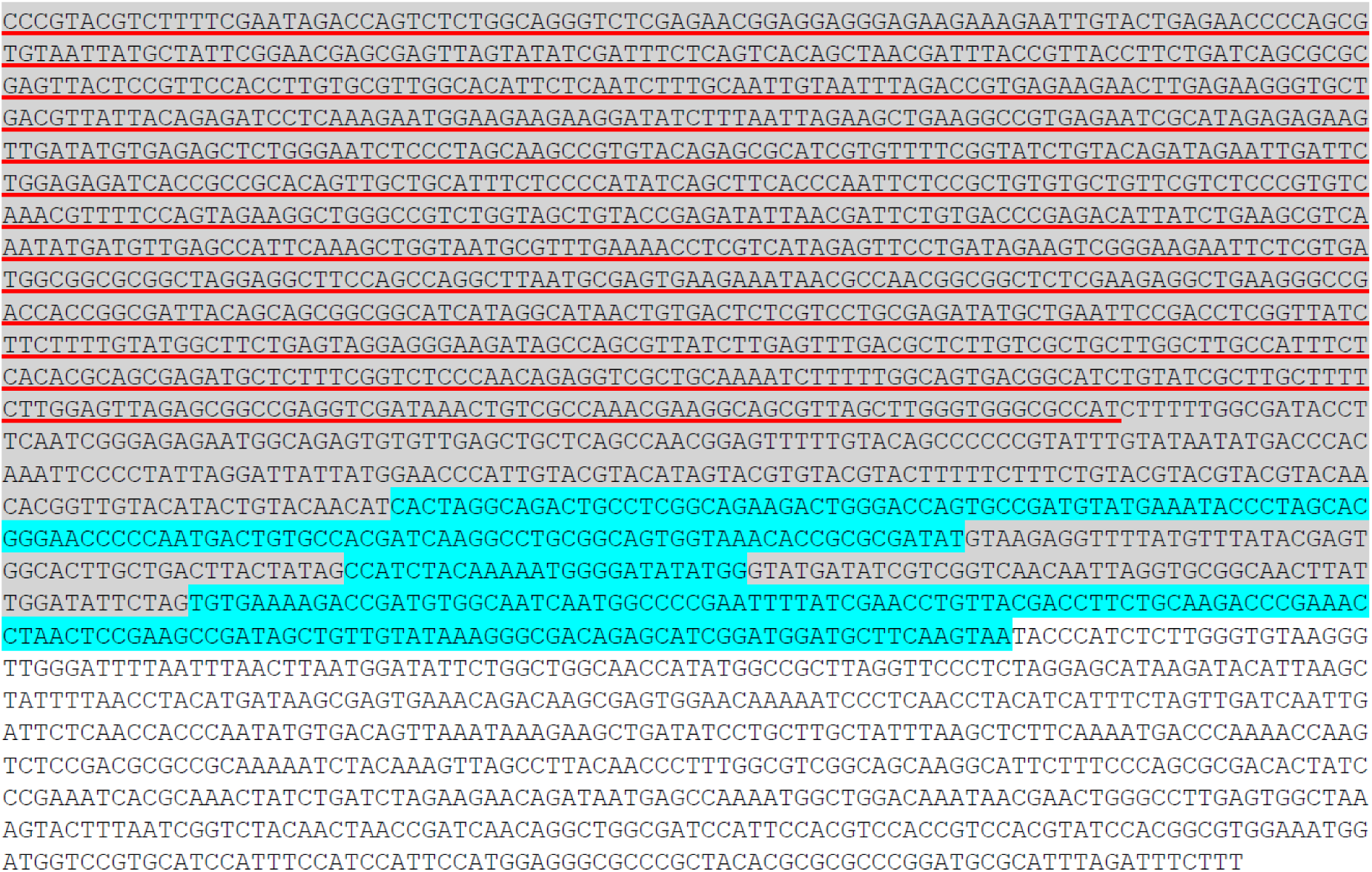
Nucleotide sequence of the *Fol*4287 *SIX8*-*PSE1* locus with TE inserted. Yellow and vivid blue highlights indicate the *SIX8b* and *PSE1* coding sequences (CDS) in opposite transcriptional orientations. Intron sequences are highlighted in gray. Red underline represents the sequence encoding a putative transposase gene.

**Supplementary Figure 8.**
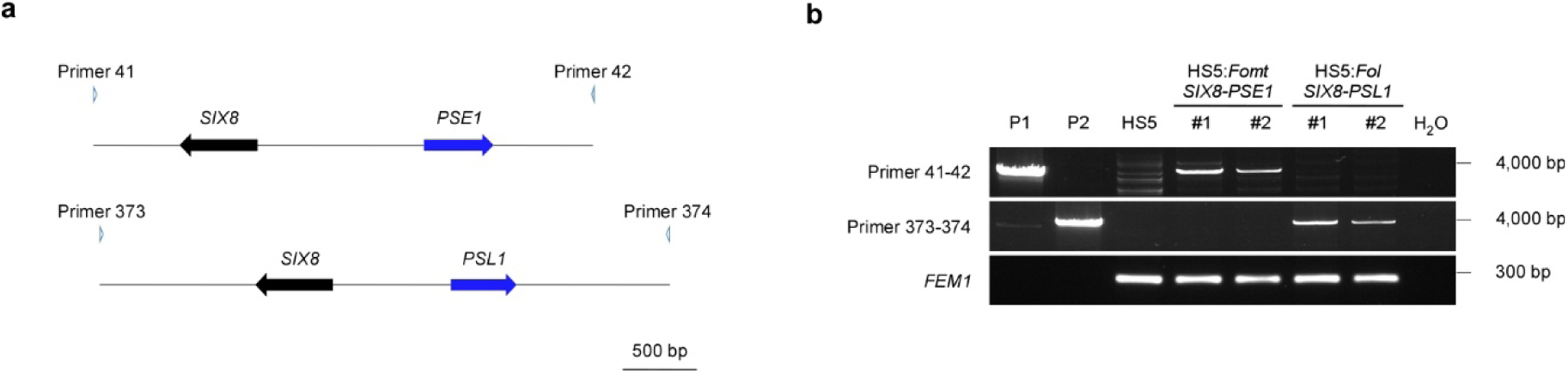
Introduction of the *Fomt SIX8*-*PSE1* locus and the *Fol SIX8-PSL1* locus into *Focn*Cong:1-1 HS5. **a,** Schematic representation of the *Fomt SIX8*-*PSE1* locus (upper) and the *Fol SIX8*-*PSL1* locus (lower). Arrowheads indicate the primer locations for PCR verification of transformation. **b,** Confirmation of *Focn*Cong:1-1 HS5 transformants introduced with the *Fomt SIX8*-*PSE1* locus (HS5:*Fomt SIX8*-*PSE1*) or the *Fol SIX8*-*PSL1* locus (HS5:*Fol SIX8*-*PSL1*) by PCR. *FEM1* was amplified as a control. Plasmid DNA containing the *Fomt SIX8*-*PSE1* locus (P1) or the *Fol SIX8*-*PSL1* locus (P2) and DNAs of HS5, two independent HS5:*Fomt SIX8*-*PSE1* strains (#1, #2) and HS5:*Fol SIX8*-*PSL1* strains (#1, #2) were used as templates.

**Supplementary Figure 9.**
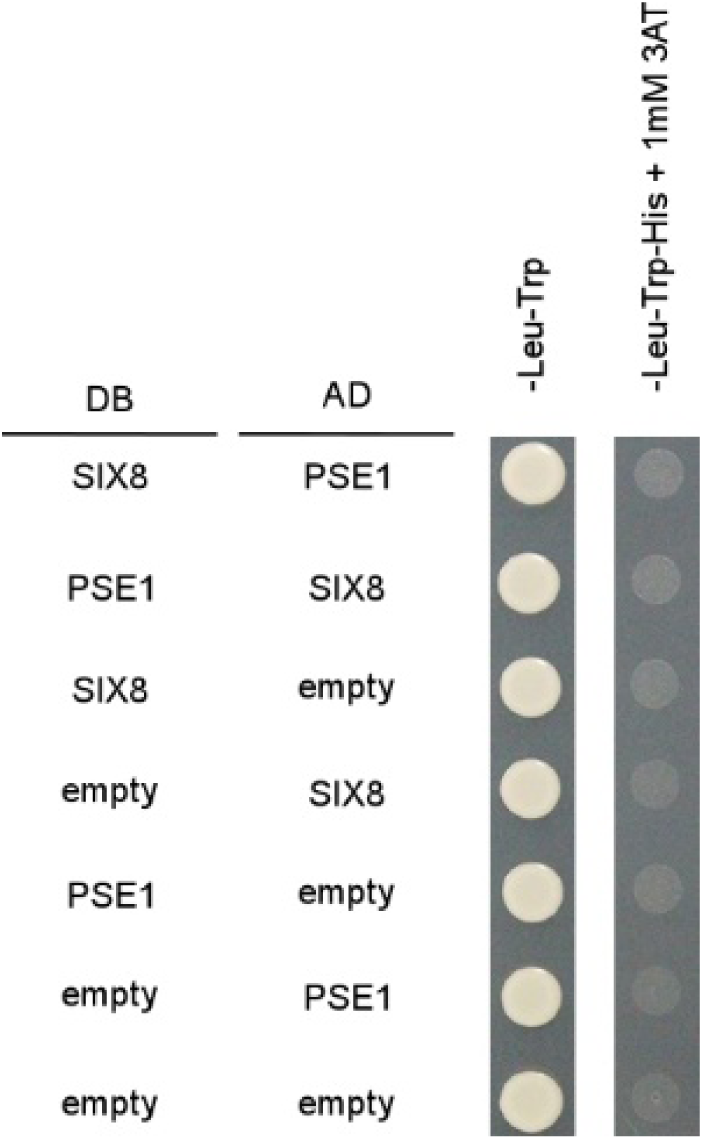
Yeast two-hybrid assay indicates no direct interaction between SIX8 and PSE1. *SIX8* and *PSE1* were cloned into bait vector (pDEST-DB; DB) and prey vector (pDEST-AD; AD). DB and AD were transformed into yeast strain *MAT*α Y8930 and *MAT*a Y8800, respectively. After mating of the yeast transformants, yeast suspension was spotted on synthetic defined (SD) medium lacking leucine and tryptophan (-Leu-Trp) for selection of diploid cells (left) and on SD medium lacking leucine, tryptophan and histidine and supplemented with 1 mM 3-amino-1,2,4-triazole (-Leu-Trp-His + 1mM 3AT) for verifying a protein-protein interaction (right).

**Supplementary Table 1.** Information for genome assembly of *F. oxysporum* isolates.

| Organism | Isolate | No. of scaffolds/contigs | GenBank assembly accession | Reference |
| --- | --- | --- | --- | --- |
| *F. oxysporum* f. sp. *conglutinans* | Cong:1-1 | 22 + 125<sup>a</sup> | in process | this study |
| *F. oxysporum* | Fo5176 | 19 | GCA_000222805.1 | Fokkens et al. 2020 |
| *F. oxysporum* f. sp. *matthiolae* | PHW726 | 66 + 517<sup>b</sup> | GCA_009755825.1 | Yu et al. 2020 |
| *F. oxysporum* f. sp. *lycopersici* | 4287 | 15 + 73<sup>c</sup> | GCA_000149955.2 | Ma et al. 2010 |
| *F. oxysporum* f. sp. *cubense* | TR4 | 15 | GCA_007994515.1 | Warmington et al. 2019 |
| *F. oxysporum* f. sp. *cubense* | 160527 | 12 | GCA_005930515.1 | Asai et al. 2019 |
| *F. oxysporum* f. sp. *radicis-cucumerinum* | Forc 016 | 33 | GCA_001702695.2 | van Dam et al. 2017 |
| *F. oxysporum* f. sp. *melonis* | 26406 | 96 | GCA_002318975.1 | van Dam et al. 2017 |
| *F. oxysporum* f. sp. *cepae* | FoC_Fus2 | 34 | GCA_003615085.1 | Armitage et al. 2018 |
| *F. oxysporum* f. sp. *vasinfectum* | TF1 | 17 | GCA_009602505.1 | Seo et al. 2020 |

<sup>a</sup>Number of scaffolds + non-scaffolded contigs.<sup>b</sup>Number of core scaffolds + scaffolds.<sup>c</sup>Number of chromosomes + non-scaffolded contigs.

**Supplementary Table 2.** *F. oxysporum* strains used in this study.

| Strain | Genotype | Reference |
| --- | --- | --- |
| *Focn* Cong:1-1 | wild type | Kawabe et al. 2005 |
| ΔSIX4 | ΔSIX4::hph | Kashiwa et al. 2013 |
| HS1 | ΔSC9 | this study |
| HS2 | ΔSC3/SC9 | this study |
| HS3 | ΔSC3/SC9/chr<sup>SC10/SC20</sup> | this study |
| HS4 | ΔSC3/SC9/chr<sup>SC10/SC20</sup>/chr<sup>SC16/SC18</sup> | this study |
| HS5 | ΔSC9/chr<sup>SC10/SC20</sup> | this study |
| HS6 | ΔSC5/SC8/SC9/chr<sup>SC10/SC20</sup> | this study |
| HS6-BLE | ΔSC5/SC8/SC9/chr<sup>SC10/SC20</sup>::ble | this study |
| HCT1 | ΔSC5/SC8/SC9/chr<sup>SC10/SC20</sup>::SC8/SC9/chr<sup>SC10/SC20</sup>/hph/ble | this study |
| HCT2 | ΔSC5/SC8/SC9/chr<sup>SC10/SC20</sup>::SC8/SC9/hph/ble | this study |
| HCT3 | ΔSC5/SC8/SC9/chr<sup>SC10/SC20</sup>::SC8/SC9/hph/ble | this study |
| HCT4 | ΔSC5/SC8/SC9/chr<sup>SC10/SC20</sup>::SC8/SC9/hph/ble | this study |
| ΔSIX4-GFP | ΔSIX4::hph/nptII/GFP | this study |
| HS2-GFP | ΔSC3/SC9::hph/GFP | this study |
| HS5-GFP | ΔSC9/chr<sup>SC10/SC20</sup>::hph/GFP | this study |
| HS5:SIX8-PSE1#1 | ΔSC9/chr<sup>SC10/SC20</sup>::SIX8-PSE1 locus/hph | this study |
| HS5:SIX8-PSE1#2 | ΔSC9/chr<sup>SC10/SC20</sup>::SIX8-PSE1 locus/hph | this study |
| HS5:SIX8#1 | ΔSC9/chr<sup>SC10/SC20</sup>::SIX8/hph | this study |
| HS5:SIX8#2 | ΔSC9/chr<sup>SC10/SC20</sup>::SIX8/hph | this study |
| HS5:PSE1#1 | ΔSC9/chr<sup>SC10/SC20</sup>::PSE1/hph | this study |
| HS5:PSE1#2 | ΔSC9/chr<sup>SC10/SC20</sup>::PSE1/hph | this study |
| ΔSIX8-PSE1#1 | ΔSIX8-PSE1 locus::hph | this study |
| ΔSIX8-PSE1#2 | ΔSIX8-PSE1 locus::hph | this study |
| ΔSIX8-PSE1#3 | ΔSIX8-PSE1 locus::hph | this study |
| ect | hph | this study |
| HS5:FomtSIX8-PSE1#1 | ΔSC9/chr<sup>SC10/SC20</sup>::FomtSIX8-PSE1 locus/hph | this study |
| HS5:FomtSIX8-PSE1#2 | ΔSC9/chr<sup>SC10/SC20</sup>::FomtSIX8-PSE1 locus/hph | this study |
| HS5:FolSIX8-PSE1#1 | ΔSC9/chr<sup>SC10/SC20</sup>::FolSIX8-PSL1 locus/hph | this study |
| HS5:FolSIX8-PSE1#2 | ΔSC9/chr<sup>SC10/SC20</sup>::FolSIX8-PSL1 locus/hph | this study |
| HS5:empty | ΔSC9/chr<sup>SC10/SC20</sup>::hph | this study |

**Supplementary Table 3.**
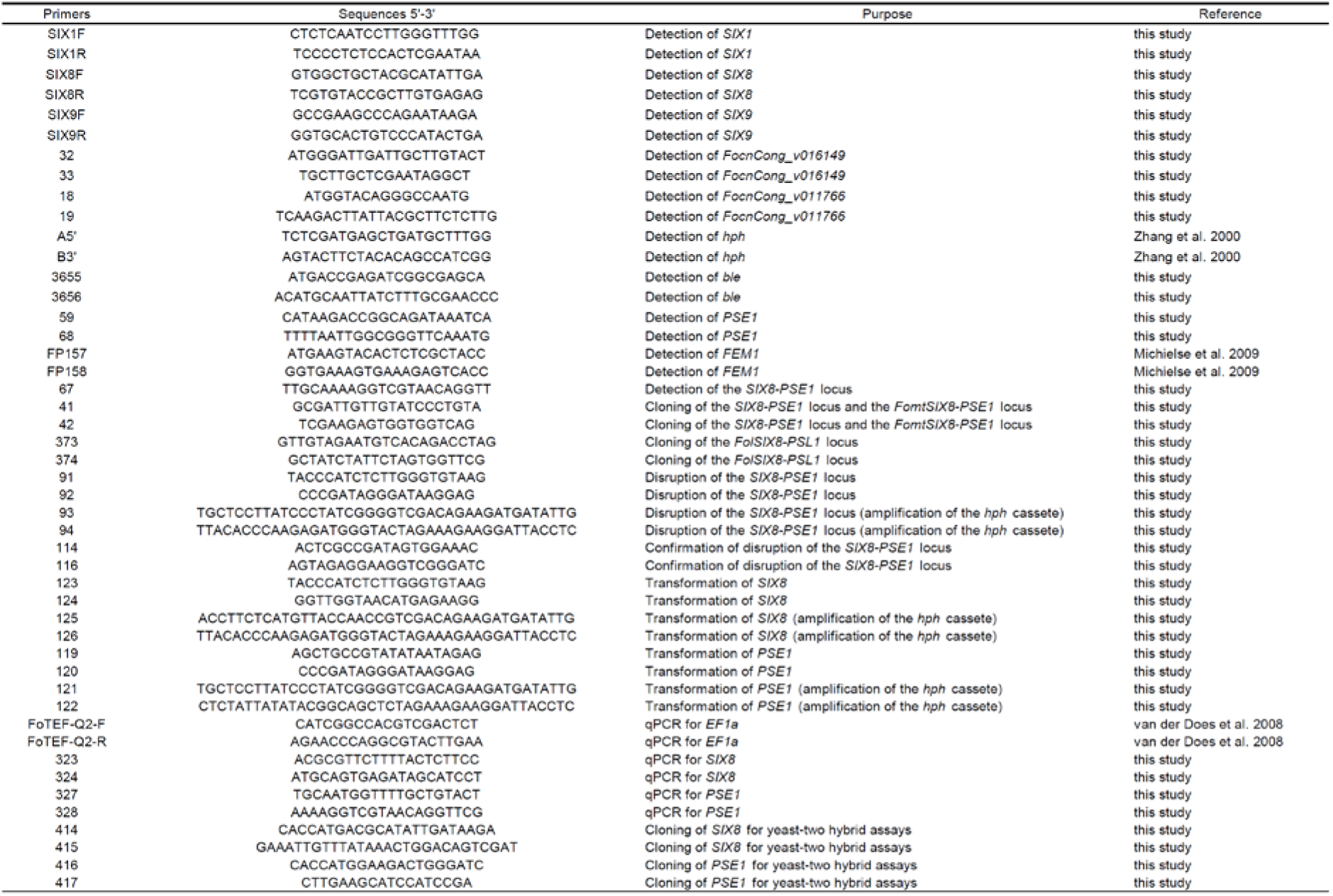
Primers used in this study.

**Supplementary Table 4.** Plasmid DNAs used in this study.

| Plasmids | Description | Resistance | Reference |
| --- | --- | --- | --- |
| pCSN43 | vector for expression of *hph* | ampicillin, hygromycin B | Staben et al. 1989 |
| pMK412 | vector for expression of *hph* and *GFP* | ampicillin, hygromycin B | Watanabe et al. 2007 |
| pMD-GEN | vector for expression of *nptII* and *GFP* | ampicillin, geneticin | Saito et al. submitted |
| pSIX8-PSE1 | vector for expression of *SIX8* and *PSE1* | spectinomycin | this study |
| pSIX8 | vector for expression of *SIX8* and *hph* | spectinomycin | this study |
| pPSE1 | vector for expression of *PSE1* and *hph* | spectinomycin, hygromycin B | this study |
| pKOSIX8-PSE1 | vector for disruption of the *SIX8-PSE1* locus | spectinomycin, hygromycin B | this study |
| pFomtSIX8-PSE1 | vector for expression of *FomtSIX8* and *FomtPSE1* | spectinomycin | this study |
| pFolSIX8-PSL1 | vector for expression of *FolSIX8* and *PSL1* | spectinomycin | this study |
| pSIX8-AD | vector for expression of *SIX8* fused with an activating domain of *GAL4* | carbendazim | this study |
| pPSE1-AD | vector for expression of *PSE1* fused with an activating domain of *GAL4* | carbendazim | this study |
| pSIX8-DB | vector for expression of *SIX8* fused with a DNA binding domain of *GAL4* | carbendazim | this study |
| pPSE1-DB | vector for expression of *PSE1* fused with a DNA binding domain of *GAL4* | carbendazim | this study |
| pDEST-AD | vector for expression of an activating domain of *GAL4* | carbendazim | Ahmed et al. 2018 |
| pDEST-DB | vector for expression of a DNA binding domain of *GAL4* | carbendazim | Ahmed et al. 2018 |

**Supplementary Table 5.** Genome sequences used for repeat element prediction.

| Organism | Isolate | N50 contig length (Mb) | GenBank assembly accession | Reference |
| --- | --- | --- | --- | --- |
| *F. oxysporum* f. sp. *cubense* | 160527 | 4.9 | GCA_005930515.1 | Asai et al. 2019 |
| *F. oxysporum* f. sp. *conglutinans* | Cong:1-1 | 4.9 | in process | This study |
| *F. oxysporum* f. sp. *cubense* | TR4 | 4.5 | GCA_007994515.1 | Warmington et al. 2019 |
| *F. oxysporum* f. sp. *radicis-cucumerinum* | Forc016 | 4.5 | GCA_001702695.2 | van Dam et al. 2017 |
| *F. oxysporum* f. sp. *lycopersici* | race 3 | 4.4 | GCA_003977725.1 | Henry et al. 2019 |
| *F. oxysporum* f. sp. *melonis* | 26406 | 4.4 | GCA_002318975.1 | van Dam et al. 2017 |
| *F. oxysporum* f. sp. *cepae* | FoC_Fus2 | 4.1 | GCA_003615085.1 | Armitage et al. 2018 |
| *F. oxysporum* f. sp. *lycopersici* | 4287 | 1.3 | GCA_003315725.1 | Ayhan et al. 2018 |

